# Causal role of the individual alpha phase in multisensory perception

**DOI:** 10.1101/2025.11.11.687884

**Authors:** Tim Rohe, Daniel Senkowski

## Abstract

The phase of alpha oscillations is proposed to reflect perceptual cycles. Yet, human EEG/MEG studies have yielded inconsistent results, perhaps because they used correlative approaches that did not consider causal phase-perception relationships. In a multi-day computational EEG study, we developed the neurobehavioral link function (NBLF) as a new approach to model individual sinusoidal relationships between the alpha phase and multisensory perception and to test for causality. Rhythmic 10 Hz visual stimulation entrained the alpha phase at 0° or 180° relative to audiovisual target onset. Entrainment effects on multisensory integration only emerged when individual NBLFs, which were related to alpha phase coupling between early audio-visual cortices, were included in the analysis. A follow-up psychophysical experiment confirmed individual phase-dependent effects at 5 and 10 Hz, but not 15 Hz. Our results suggest that the individual alpha phase causally modulates perceptual cycles in multisensory integration and advocate for personalized, phase-targeting strategies in neurotechnologies.

## Introduction

Alpha-band (8-12 Hz) oscillations have been found to support both bottom-up^1,2^ and top-down^3,4^ processing by modulating the flow of sensory information in the brain^5,6^. The waxing and waning of alpha oscillations presumably reflects periodic alterations in neural excitability^1,7,8^, providing temporal windows for attentional sampling and perceptual integration^9–13^. However, human electroencephalography (EEG) and magnetoencephalography (MEG) studies investigating the phase of alpha oscillations and their influence on perception have produced conflicting results, yielding both positive^14–20^ and negative^21–26^ outcomes. The factors contributing to these conflicting results are not well understood.

One possible explanation for this heterogeneity is that the relationship between alpha phase and perception has primarily been investigated with correlational approaches^27^. These approaches are limited because they are susceptible to false positive or negative findings due to mediating or confounding variables, such as other neurophysiological mechanisms^28^ or attention^9,29^. Furthermore, the relationship between the alpha phase and perception can vary significantly between observers^30^, as shown in the auditory^31,32^, visual^14,33,34^, somatosensory^35^ and audiovisual^19^ modalities. The same alpha phase may correspond to high perceptual performance in one observer but low performance in another. Thus, when averaging across observers, differential relationships between the alpha phase and perception effects can cancel each other out^30^. To account for this, some studies have used relative phase opposition measures, in which trials are sorted according to their behavioural or perceptual outcome for each observer^14,17,30,36^. While these studies better account for individual phase-perception relationships, they are also limited by their correlative nature. Therefore, the mixed findings regarding the effects of the alpha phase on perception may be attributed to the use of correlative approaches, as well as to interindividual differences in the relationship between phase and perception. Only studies that manipulate the alpha phase directly can establish a causal link between phase and perception, but these studies also need to consider individual phase-perception relationships.

Several studies using sensory stimulation^37–42^ or transcranial alternating current stimulation (tACS)^43–45^ have examined the effects of alpha phase manipulation on perception. These studies also produced conflicting results^46^. While some studies reported no perceptual effects^40,41,45^, others found that rhythmic sensory or electrical stimulation can entrain the EEG/MEG alpha phase and, in parallel, influence perception^37–39,42–44^. However, these studies did not demonstrate that the perceptual effect was directly caused by the entrained alpha phase. Non-specific mechanisms, such as the entrainment of higher-order harmonics^47^, remote network effects^48^ or transcutaneous stimulation in tACS^49^, could have mediated the effects observed in these studies^50^. Demonstrating that the entrained alpha phase directly affects perception remains an important, yet under-explored, path to establishing causality.

To address this missing link, we developed a novel modelling approach to study direct causal relationships between the alpha phase and perception (Fig. 1). We refer to this approach as the neurobehavioural link function (NBLF). The NBLF describes the strength of the phase-perception modulation (i.e. its amplitude) and the phase of the alpha cycle during which the perceptual function is high or low (i.e. its shape). In a multi-day computational EEG study, we investigated the causal role of the alpha phase in audiovisual integration by using the NBLFs to predict individual alpha phase entrainment effects on multisensory perception. We adapted an established multisensory paradigm^19,51^ (Fig. 1A-B) and modelled NBLFs as individual sinusoidal relationships between the alpha phase and the multisensory *causal prior*^52,53^, reflecting a priori multisensory integration tendencies (Fig. 1C). The critical experimental manipulation involved entrainment of the endogenous alpha phase using sinusoidal visual stimulation at 10 Hz, at either on-phase (0°) or off-phase (180°) relative to the onset of audiovisual target stimuli. This design enabled us to examine the direct causal pathway from the entrainment stimulus to modulated alpha phase to perceptual outcomes (Fig. 1D). We tested three scenarios for the effects of the entrained alpha phase on multisensory perception (Fig. 1E): (i) no causal effect, (ii) a causal effect with low interindividual variability in phase-perception relationships, and (iii) a causal effect with high interindividual variability in these relationships. Our results, corroborated by a follow-up psychophysical experiment, strongly favour the third scenario.

**Figure 1.**
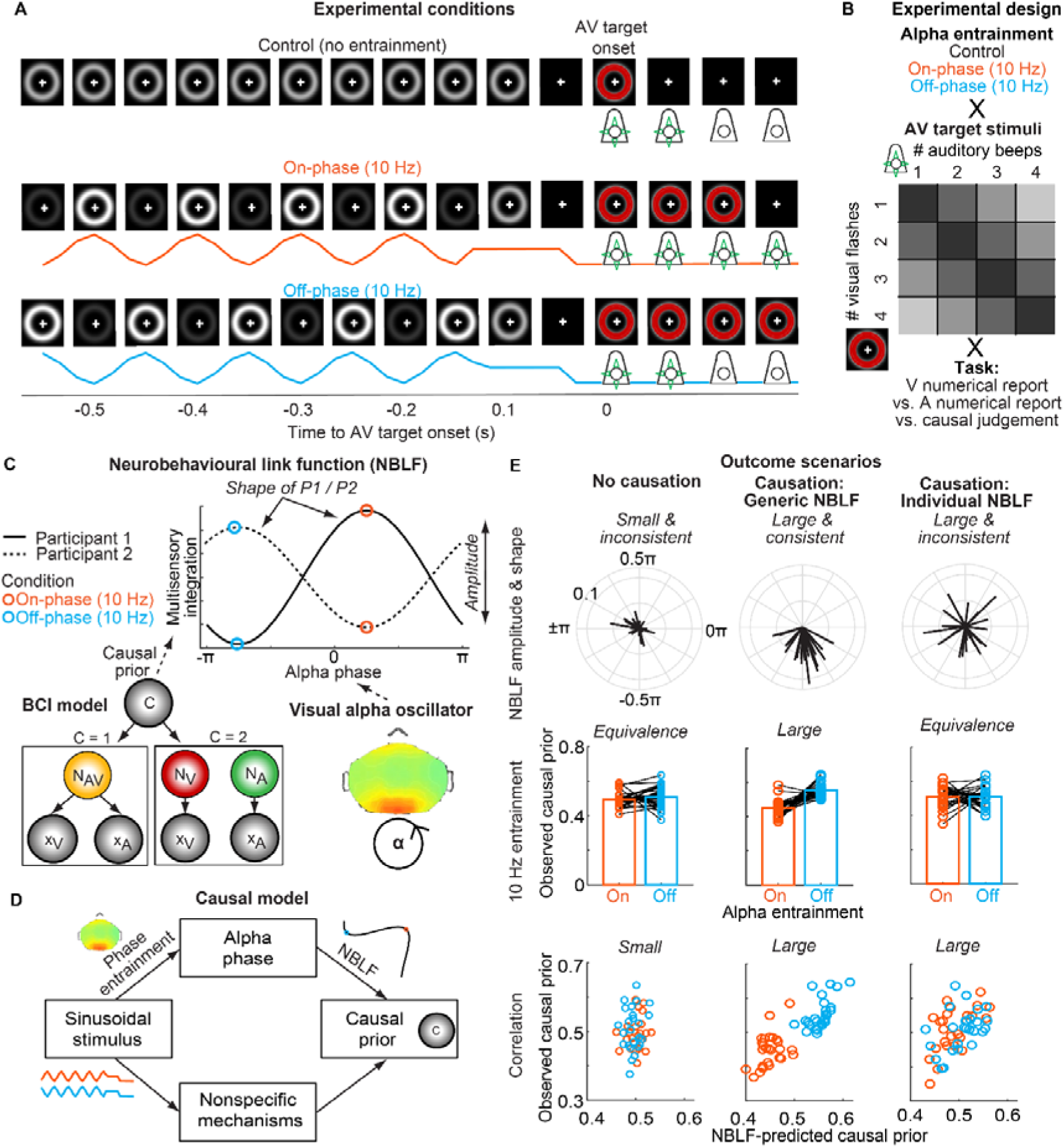
Experimental setup, graphical representation of the neurobehavioural link function, and causal model with outcome scenarios. **(A)** The EEG experiment consisted of a non-entrainment control condition, in which a static circle was presented on a screen, and two experimental conditions (10 Hz on-phase or off-phase sinusoidal visual stimulation) to entrain endogenous alpha oscillations prior to presenting AV target stimuli. The on-phase and off-phase of the entraining stimuli were defined relative (0° or 180°, respectively) to the AV target stimulus onset. In all conditions, a grey circle of the same luminance was presented 100 ms before the target onset. **(B)** Audiovisual stimuli had different numeric disparities (from 0, black, to 3, light-grey). Participants reported the number of visual or auditory stimuli, or judged the causal structure of the AV targets. **(C)** The illustration shows the NBLF for two simulated participants with opposite alpha phase-perception relationships. The NBLF models the unique sinusoidal relationship between an individual’s visual alpha phase and multisensory integration as measured by the BCI model’s multisensory causal prior. Depending on the amplitude and shape of the NBLF, the causal prior rises and falls over the cycle of the alpha oscillator. When two participants have different preferred and non-preferred phases (i.e. different NBLF shapes) and the alpha phase is entrained at opposite phases (i.e. on-phase versus off-phase), the individual NBLFs predict opposite entrainment effects on the causal prior. **(D)** The causal model posits that the sinusoidal visual stimulation at 10 Hz entrains the alpha phase, which then modulates the causal prior according to a generic or individual NBLF. Crucially, investigating this causal pathway bypass the potential mediating effect of nonspecific neural mechanisms. **(E)** This illustration shows three possible outcome scenarios for the effect of entrained alpha phase on the causal prior, using simulated data. *Left column*: The alpha phase has no causal effect. In this case, the NBLF amplitudes would be small and inconsistent across observers, and the shapes are uniformly distributed around the circle. Under entrainment conditions, equivalent causal priors are expected at the group level for on-phase and off-phase entrainment, and the predictions for the causal prior from the fitted individual NBLFs and the empirically observed causal priors are not correlated. *Middle column*: The alpha phase has a causal effect according to a generic NBLF. In this case, the NBLFs would have large amplitudes and consistent shapes across observers, i.e. the shapes cluster around a particular angle. This would lead to significant differences in causal priors between entrainment conditions and result in a significant correlation between the predictions for the causal prior from the fitted NBLFs and the observed causal priors. *Right column*: The alpha phase has a causal effect according to individual NBLFs. In this case, the NBLF amplitudes would be large but the shapes would be inconsistent across observers. In contrast to the generic NBLF scenario, equivalent causal priors are expected at the group level for the two entrainment conditions, because averaging across participants cancels the sinusoidal modulation of the causal prior. Yet, a significant correlation between the predictions for the causal prior from the fitted individual NBLFs and the observed causal priors demonstrates the causal effect of the individual alpha phase on multisensory integration.

## Results

### Experimental setup and analysis approach

To study the effects of the alpha phase on multisensory integration, we presented ∼3300 trials to each of 26 human observers, collected across two EEG recording sessions on different days. In the control condition, trials began with a static visual stimulus. In two different entrainment conditions, trials began with visual stimuli that flickered sinusoidally at 10 Hz with on-phase (0°) or off-phase (180°)^41,54^ relative to the onset of audiovisual (AV) flash-beep target stimuli (Fig. 1A-B)^55–57^. In all three conditions, the same grey circle was presented between -166 ms and -50 ms before the AV target onset, so that the offset responses were identical for all conditions. To measure observers’ multisensory integration, participants reported the number of visual flashes or auditory beeps in visual and auditory numerical report task, respectively. Multisensory integration results in crossmodal biases (CMBs)^58,59^ of visual signals on auditory reports and of auditory signals on visual reports. Because observers integrate multisensory signals only if they infer a common cause^52,53,58^, participants judged whether they perceived a common (i.e. auditory and visual inputs have the same source) or independent causes (i.e. auditory and visual input have different sources) of the flash-beep stimuli in a causal judgements task.

To test the causal model (Fig. 1D) with the three outcome scenarios (Fig. 1E), we first examined the effects of entrainment on multisensory integration as measured by the behavioural indices of CMB and causal judgements. We used a computational Bayesian causal inference (BCI) model^52,53^ to quantify the a priori tendency of observers to integrate multisensory information, i.e. the model’s multisensory *causal prior* parameter. Next, we assessed how the flickering visual stimuli entrained the phase of an alpha oscillator in the visual cortex. Finally, we modelled NBLFs in the non-entrainment control condition (Fig. 1C) and used these NBLFs to predict the individual effects of 10 Hz visual entrainment on the multisensory causal prior.

### Behavioural indices of multisensory integration show no differences between on-phase and off-phase entrainment conditions

At the behavioural level, we first analysed the CMB (Fig. 2A) and causal judgements (Fig. 2B) to determine whether any characteristics of multisensory integration were present and whether the entrainment stimuli influenced multisensory integration. When a small disparity between the number of flashes and beeps indicated a common cause, the number of beeps had the greatest crossmodal influence on visual reports, and the number of flashes biased the auditory reports (i.e. interaction between task relevance x numeric disparity, F_1.6, 38.8_ = 132.244, p < 0.001, part. η^2^ = 0.841, BF_10_ > 100; see Fig. 2A and supplemental results with Tab. S1, Figs. S1 and S2). A small numeric disparity also increased the proportion of common-cause judgements (F_1.5,37.9_ = 165.466, p < 0.001, part. η^2^ = 0.869, BF_10_ > 100; Fig. 2B and Tab. S1). Furthermore, there was no difference in the key profiles of CMB and causal judgements between on-phase and off-phase entrainment conditions (CMB: F_1,25_ = 0.250, p = 0.621, part. η^2^ = 0.010, BF_10_ = 0.149; causal judgements: F_1, 25_ = 0.001, p = 0.970, part. η^2^ < 0.001, BF_10_ = 0.141; Tab. S1). Thus, both CMB and causal judgements exhibited the defining characteristics of multisensory integration when performing causal inference^19,58,60^. Additionally, there were no significant differences in these indices between the on-phase and off-phase entrainment conditions.

**Figure 2.**
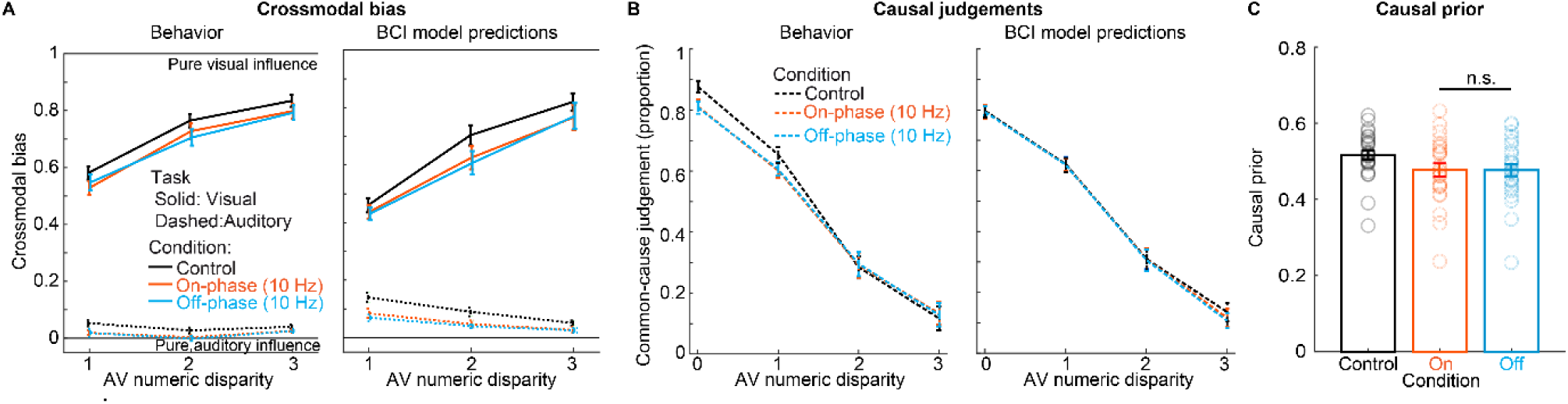
Behavioural data show no differences in multisensory integration between on-phase and off-phase entrainment conditions. **(A)** Crossmodal bias (CMB; mean across participants ± SEM; n = 26) was computed from participants’ visual and auditory numerical reports (*left*) and the BCI model predictions (*right*) and plotted for the three experimental conditions. CMB defines the relative influence of the audiovisual signals on the numerical reports and ranges from pure auditory influence (CMB = 0) to pure visual influence (CMB = 1). The key profile of CMB was equivalent between the two entrainment conditions. **(B)** Causal judgements computed from participants’ behaviour (*left*) and the BCI model’s predictions (*right*), are depicted as in (A). Causal judgements decreased with increasing numeric disparity but did not differ between the on-phase and off-phase conditions. **(C)** The causal prior parameter of the BCI model was around 0.5 across all conditions. This indicates that observers considered the possibility of one common source or two independent sources of the AV target stimuli to be equally likely a priori. Critically, the causal prior did not differ between the on-phase and off-phase entrainment conditions (Tab. S3).

Next, we fitted the BCI model to the behavioural data of the participants and compared the multisensory causal prior between entrainment conditions. The BCI model outperformed five competing computational models in the model comparison (supplemental results and Tab. S2). It also accurately predicted the participants’ CMB and causal judgement profiles (Fig. 2B; Fig. S1). Interestingly, there was no significant difference in the causal prior between the two entrainment conditions (t_25_ = 0.140, p = 0.889, Cohen’d = 0.027, BF_10_ = 0.209; Fig. 2C; Tab. S3). Furthermore, there were no significant differences between the two entrainment conditions in the other BCI model parameters (all p_cor_ > 0.05; see supplemental results, Tab. S3, Fig. S3). Taken together, the analysis of behavioural data revealed equivalent multisensory integration effects for on-phase and off-phase entrainment conditions. This lack of entrainment effects provides evidence against a generic NBLF (Fig. 1E, *middle*), but is consistent with the no causation (Fig. 1E, *left*) and the causation by individual NBLFs (Fig. 1E, *right*) scenarios.

### Rhythmic 10 Hz visual stimulation entrains ongoing alpha oscillations in early visual areas

In this analysis, we examined whether the on-phase and off-phase visual stimulation entrained an endogenous alpha oscillator with opposite phases in the early visual cortex^39,47,54^. Evidence of alpha phase entrainment is necessary to prove its causal role in multisensory integration (Fig. 1D). Our analysis of event-related potentials (ERPs) in the pre-target period (i.e. -1 to 0 seconds) and locked to the onset of AV target stimuli revealed that the rhythmic visual stimulation robustly evoked 10 Hz activity over the visual cortex (Fig. 3A). Mean phase angles were opposite for on-phase versus off-phase entrainment and this activity could be attributed to cortical sources in the early visual cortex (Fig. 3B). Specifically, the phase of the entrained 10 Hz oscillations significantly deviated from a uniform circular distribution, with highly consistent phase differences of 180° between the entrainment conditions. To examine the 10 Hz phase more closely, we calculated the intra-trial phase coherence (ITPC). For the two entrainment conditions, but not the control condition, the ITPC at 10 Hz was higher than the baseline in a cluster that began 750 ms before the AV target onset (Fig. 3C). As expected for rhythmic stimulation in a nonlinear system, evoked activity was also present at higher-order harmonics at 20, 30, and 40 Hz^47,61–63^. Interestingly, the time courses of the occipital 10 Hz phase distributions gradually and smoothly followed the on–off phase of the rhythmic stimuli after their onset (Fig. 3D). Similar results were obtained for ITPC and phase distribution in the source-localised responses in early visual areas (see supplemental Fig. S4).

**Figure 3.**
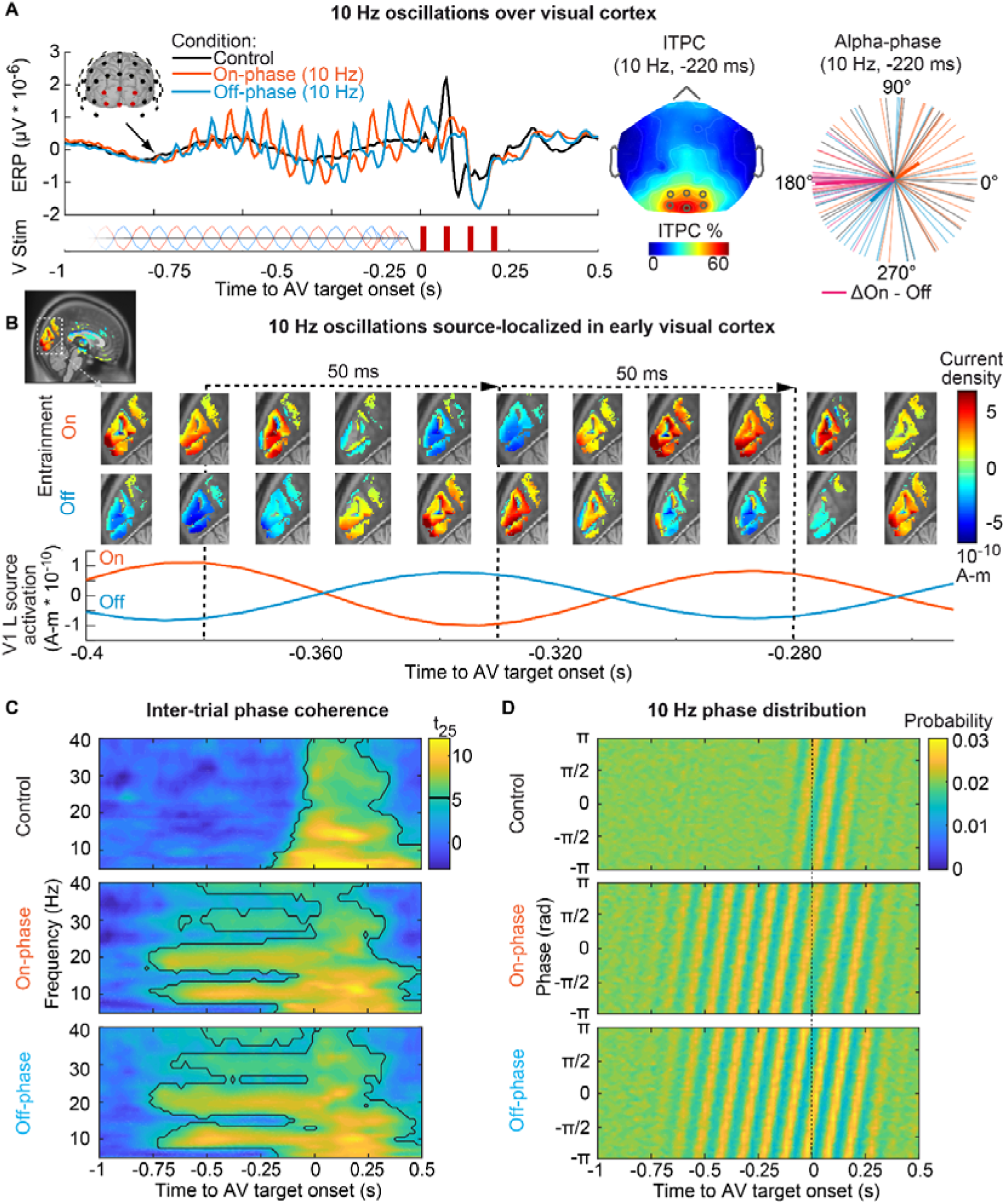
On-phase and off-phase 10 Hz visual stimulation resulted in phase-locked 10 Hz oscillations with opposite phase angles in the early visual cortex. **(A)** *Left:* Grand average ERPs at electrodes over the visual cortex locked to the onset of AV target stimuli. *Middle:* In the two entrainment conditions, average 10 Hz ITPC at -220 ms was confined to occipital electrodes. *Right:* The polar plot of the 10 Hz phases of individual participants at -220 ms shows significant phase clustering in the on-phase (Raleigh test, z = 3.382, p = 0.032, BF_10_ = 0.248) and off-phase (z = 3.942, p = 0.018, BF_10_ = 0.454) conditions. The individual circular phase difference between on-phase and off-phase entrainment clustered strongly at 180° (z = 23.979, p < 0.001, BF_10_ > 100). No phase clustering was found in the control condition (z = 0.351, p = 0.708, BF_10_ = 0.011). The thick line shows the resultant. **(B)** Source-localised bandpass-filtered (8-12 Hz) ERPs demonstrate rhythmic alpha activity in the early visual areas (V1/V2), with reversed amplitude for the on-phase and off-phase entrainment conditions. **(C)** Time-frequency t-value maps of ITPC for the three entrainment conditions. These were computed from occipital electrodes, as shown in panel A. Significant clusters (p < 0.001; two-sided cluster-based randomisation t_25_ test) are marked by a solid line. **(D)** The average distribution of 10 Hz alpha phase shows a continuous ∼180° phase shift between on-phase and off-phase stimulation. In the control condition, the phase locking is likely caused by the AV target stimuli. Notably, the 10 Hz phase entrainment in the entrainment conditions persists for several hundred milliseconds longer than in the control condition.

So far, the findings have demonstrated that the on-phase and off-phase stimulation result in robust 10 Hz oscillations of opposite phases in the visual cortex. However, it is possible that these oscillatory response patterns merely reflect an accumulation of ERPs over time due to the rhythmic 10 Hz stimulation, but without entrainment of an endogenous oscillator^64–69^. Fig. 3A shows that the 10 Hz responses in visual cortex increased gradually after the onset of the rhythmic 10 Hz visual stimulation. This suggests a neural entrainment mechanism rather than a superposition of ERPs, for which power modulations and habituation effects would be expected^62,70^. Nevertheless, we conducted further analyses to address this important issue. These analyses revealed clear evidence for the entrainment of endogenous oscillations (Fig. 4). Firstly, despite the robust 10 Hz phase locking measured by ITPC, we found no differences in total 10 Hz power between the entrainment and the control conditions (Fig. 4A). Secondly, increased power and ITPC values were evident in the entrainment conditions compared to the control condition at the second and third harmonics (i.e. 20 and 30 Hz; Fig. 4B). This may be due to nonlinear neural dynamics elicited by the interaction between an external stimulus and endogenous oscillations^61,63,71^. Thirdly, we investigated how rhythmic stimulation at 10 Hz modulated the individual alpha frequency (IAF), which was calculated from resting-state EEG data. Participants’ IAFs transiently adapted to the extraneous 10 Hz stimulation frequency before returning to baseline levels (Fig. 4C). Furthermore, although participants exhibited alpha power peaks at their preferred frequencies prior to entrainment (supplemental Fig. S5), both groups (IAF < 10 Hz and IAF ≥ 10 Hz) demonstrated consistent and robust entrainment at 10 Hz, with comparable power levels during entrainment (Fig. 4B, left panel). This transient adaptation of the alpha eigenfrequency is expected when an internal oscillator adapts to the frequency of a salient external entraining stimulus^64,72,73^. Finally, we examined 10 Hz oscillations in null-event trials, i.e. trials without AV target stimuli. In these trials, both the on-phase and off-phase stimuli entrained the phase of alpha oscillations, with a constant mean phase angle difference of 180° (Fig. 4D). Interestingly, the consistency (resultant length) of the 180° phase difference was stronger than that of the on-phase and off-phase stimulation itself. This ‘entrainment echo’^39,41,73^ persisted for ∼2 alpha cycles after the offset of the entraining stimuli. This pattern was also observed in experimental conditions involving AV target stimuli (Figs. 3A and D). In summary, these results provide strong evidence that the 10 Hz on-phase and off-phase visual stimulation entrains endogenous visual alpha oscillations at opposite phases.

**Figure 4.**
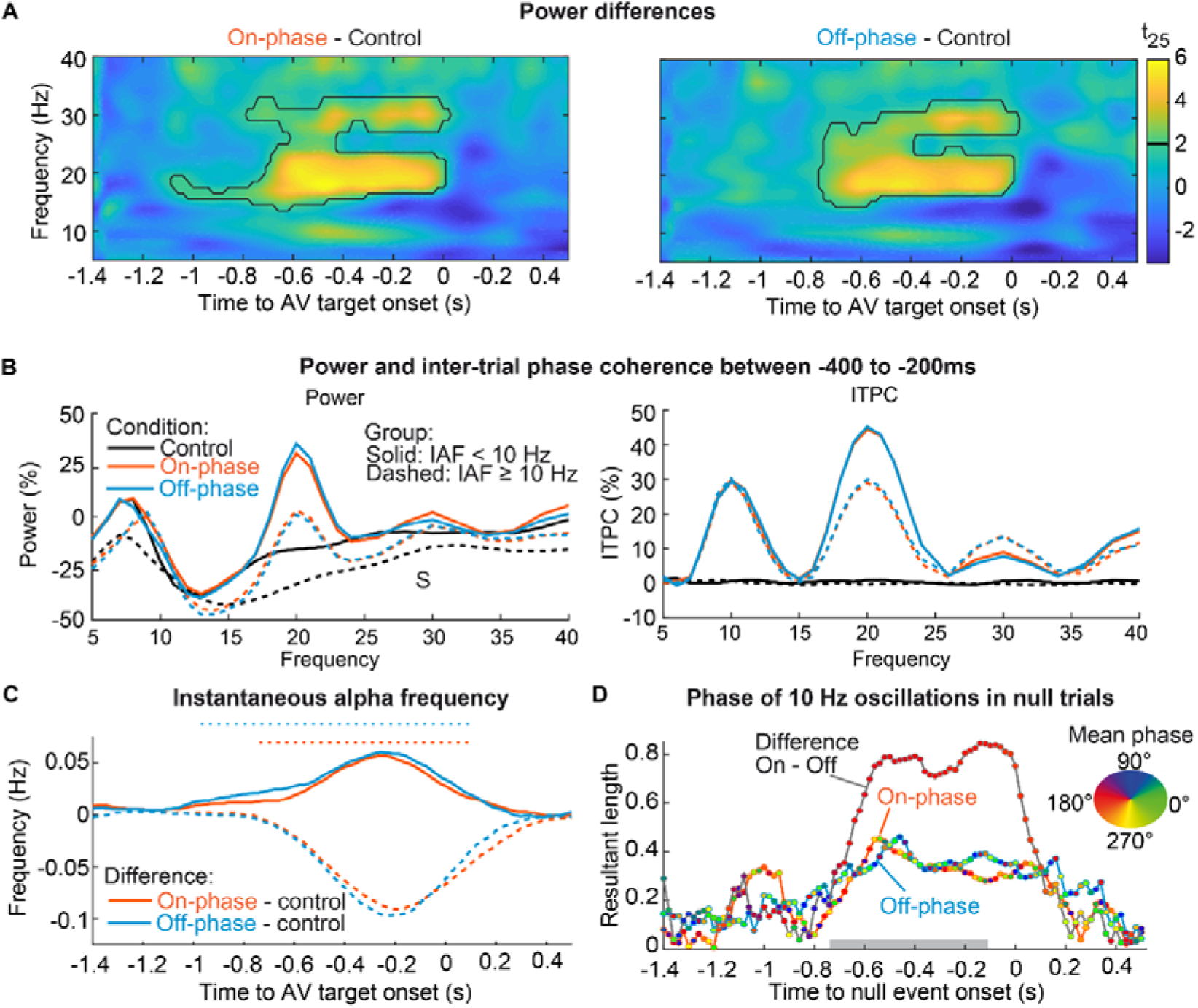
Analysis of power, ITPC, individual and instantaneous alpha frequency, and 10 Hz phase provides evidence for entrainment of endogenous alpha oscillations. **(A)** Time-frequency t-value maps of the power difference between the entrainment conditions (on-phase and off-phase) compared to the control condition. These were computed across six occipital electrodes (Fig. 3A, middle panel). Significant clusters of power differences (p ≤ 0.001, two-sided cluster-based randomisation t_25_ test) were found at the second (20 Hz) and third (30 Hz) harmonics, but not at the 10 Hz stimulation frequency. No power differences were observed between the on-phase and off-phase entrainment conditions across the 5-40 Hz power spectrum (all clusters p > 0.05). **(B)** Power (left) and ITPC (right) plotted as a function of frequency and individual alpha frequency (IAF). The data (normalised means relative to baseline) are shown for individuals with an IAF < 10 Hz (n = 10, solid line) and IAF ≥ 10 Hz (n = 16, dashed line). **(C)** The time course of the instantaneous alpha frequency is shown for the on-phase and off-phase entrainment conditions compared to the control condition for participants with IAF < 10 Hz (solid line) and IAF ≥ 10 Hz (dashed line). Color-coded horizontal dotted lines indicate clusters of significant contrasts (p < 0.01; two-sided two-sample cluster-based randomisation t_24_ test). **(D)** The time course of the 10 Hz phase angle is shown for the ‘null trials’ in the on-phase and off-phase entrainment conditions, as well as the circular difference between them. The y-axis shows the resultant length of the alpha phase across the sample. The mean phase angle of each sample is color-coded. The average duration of the entraining stimulus is indicated by the rectangular grey box. The 180° phase difference between the entrainment conditions (on-phase minus off-phase) was stronger than the phase consistency within the entrainment conditions and persisted for a few hundred milliseconds after the entrainment stimulus ceased.

### Neurobehavioural link functions show a large interindividual variability and correlate with audiovisual integration in the control condition

The next step of the analysis involved investigating whether the ongoing alpha phase correlated with multisensory integration when considering individual NBLFs. First, we calculated the NBLFs from the behavioural and EEG data of the control condition, separately for each observer (see Methods). To achieve this, we estimated the fluctuation of the causal prior as a function of the prestimulus alpha phase over the visual cortex by fitting the causal prior in the BCI model as a continuous sinusoidal function of the alpha phase. This was done across trials and at different time points, resulting in individual NBLFs (Figs. 5A-B). At each time point, the NBLF reflects the strength of the alpha phase modulation, i.e. the NBLF amplitude, as well as which phases of the alpha cycle were associated with a high or low causal prior. These shapes of the NBLFs, which characterise the individual sinusoidal relationship between the alpha phase and multisensory integration, differed significantly between observers. Uniformly distributed circular vectors (Raleigh test at -160 ms: z = 2.765; p = 0.062, BF_10_ = 0.129; see supplemental Fig. S6 for all time points), provided substantial evidence in favour of individual NBLFs (Fig. 1E, *right*). Furthermore, in agreement with our previous report^19^, a significant NBLF amplitude was found across observers from 340 ms to 180 ms before AV target onset (p < 0.05, one-sided cluster-based corrected randomisation t_25_ test; Fig. 5C). Of particular interest was the question of the neural mechanisms underlying the interindividual variability in NBLFs. To address this, we regressed the NBLF amplitude before AV target onset on prominent neural mechanisms of multisensory integration^19,74–79^, that is the occipital alpha dipole amplitude, phase coupling between primary visual areas and parietal areas, phase coupling between primary visual and primary auditory areas, and the instantaneous alpha frequency. Interestingly, the NBLF amplitude was best explained by alpha-band phase coupling between early visual and early auditory regions, and to a lesser extent by the occipital dipole amplitude (Fig. 5D). Together, these findings demonstrate a correlative relationship between the alpha phase and the multisensory causal prior when modelling individual NBLFs.

**Figure 5.**
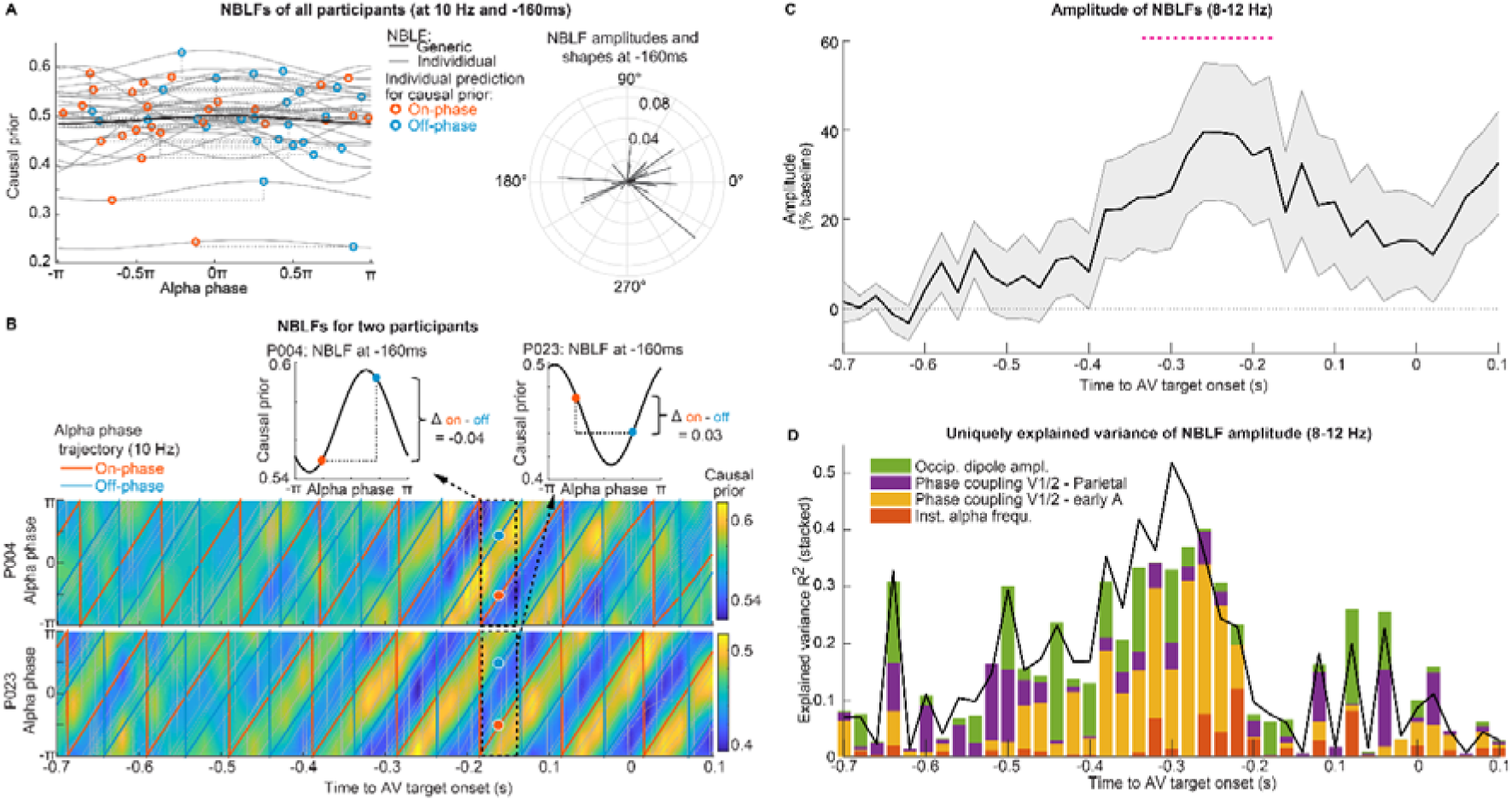
Neurobehavioural link functions, which characterise individual alpha phase-perception relationships, correlate with multisensory integration and were associated with phase coupling between early audio-visual cortices. **(A)** *Left:* Individual NBLFs were fitted to the prestimulus alpha phases in the control condition for a cluster of six electrodes located over the visual cortex (see Fig. 3A, *middle*). For each participant, the NBLFs predicted the strength of multisensory integration (i.e. the multisensory causal prior) for both the 10 Hz on-phase entrainment (red dot) and off-phase entrainment (blue dot) conditions. *Right:* The shapes of the NBLFs varied significantly across participants, as shown by their uniform distribution. **(B)** NBLFs for two participants (P004 and P023) 160 ms before the AV target stimulus onset. The entrained alpha phase trajectories in the on-phase and off-phase stimulation conditions are shown on top (blue and red lines) and show a 180° offset. The NBLFs of the two observers have opposite shapes. For these two observers, the individual NBLFs predict opposite effects of the on-phase (red dot) and off-phase (blue dot) stimulation on multisensory integration. **(C)** The amplitude of the NBLFs for 8-12 Hz oscillations increased significantly from 340 ms to 180 ms before the AV target onset. The horizontal dotted line indicates a significant cluster (p < 0.05, one-sided cluster-based randomisation t_25_ test; dotted horizontal magenta line) relative to the baseline. **(D)** Proportions of the variance in interindividual differences in NBLF amplitude that are uniquely explained by designated neural mechanisms. The black lines represent the variance jointly explained by all predictors, i.e. the full model. Within the time window of -340 ms to -180 ms, NBLF amplitudes were primarily related to alpha-band phase coupling between the early visual and early auditory cortices and to a lesser extent to the occipital dipole amplitude.

### Neurobehavioural link functions predict the individual causal effect of alpha phase entrainment on multisensory integration

Thus far, we have established two critical findings: Firstly, the on-phase and off-phase 10 Hz visual stimulation reliably entrained alpha oscillations in the early visual cortex with opposite phase angles relative to the AV target onset (Figs. 3-4). Secondly, we found that the alpha phase in the control condition was correlated with audiovisual integration when considering individual NBLFs (Fig. 5). Next, we investigated whether the entrained alpha phase drives multisensory integration in accordance with the causality criteria^50^. To demonstrate this, we had to show that the entrained alpha phase influenced multisensory perception differentially in the on-phase than in the off-phase condition when considering individual NBLFs (Fig. 1D). When we did not consider individual NBLFs, there were no differences in multisensory integration between the entrainment conditions (Fig. 2), which argued against the existence of generic NBLFs (Fig. 1E, *middle*). As illustrated in Figs. 5A and 5B, the NBLFs, which were derived solely from the control condition, enabled us to predict the individual effects of on-phase and off-phase stimulation on multisensory integration and to compare them with the observed effects. This analysis revealed positive relationships between the individual NBLF-predicted and the empirically observed multisensory causal priors for the prestimulus time interval in the alpha band (8-12 Hz). Specifically, these relationships occurred between 160 to 40 ms (mean r = 0.385; p = 0.0148) and between 360 to 300 ms (mean r = 0.378; p = 0.048) before AV target onset (Fig. 6A). The results demonstrate that the entrained visual alpha phase influences multisensory integration in a causal manner. To investigate whether these relationships could also be found at other frequencies, we analysed a broader frequency range of 5-20 Hz. This follow-up analysis revealed that the effect emerged specifically within the 8-12 Hz frequency range, from approximately 400 to 40 ms before AV target onset (Fig. 6B; p < 0.001). This also shows that the there is no effect at the second harmonic (i.e. 20 Hz) of the 10 Hz visual stimulation.

**Figure 6.**
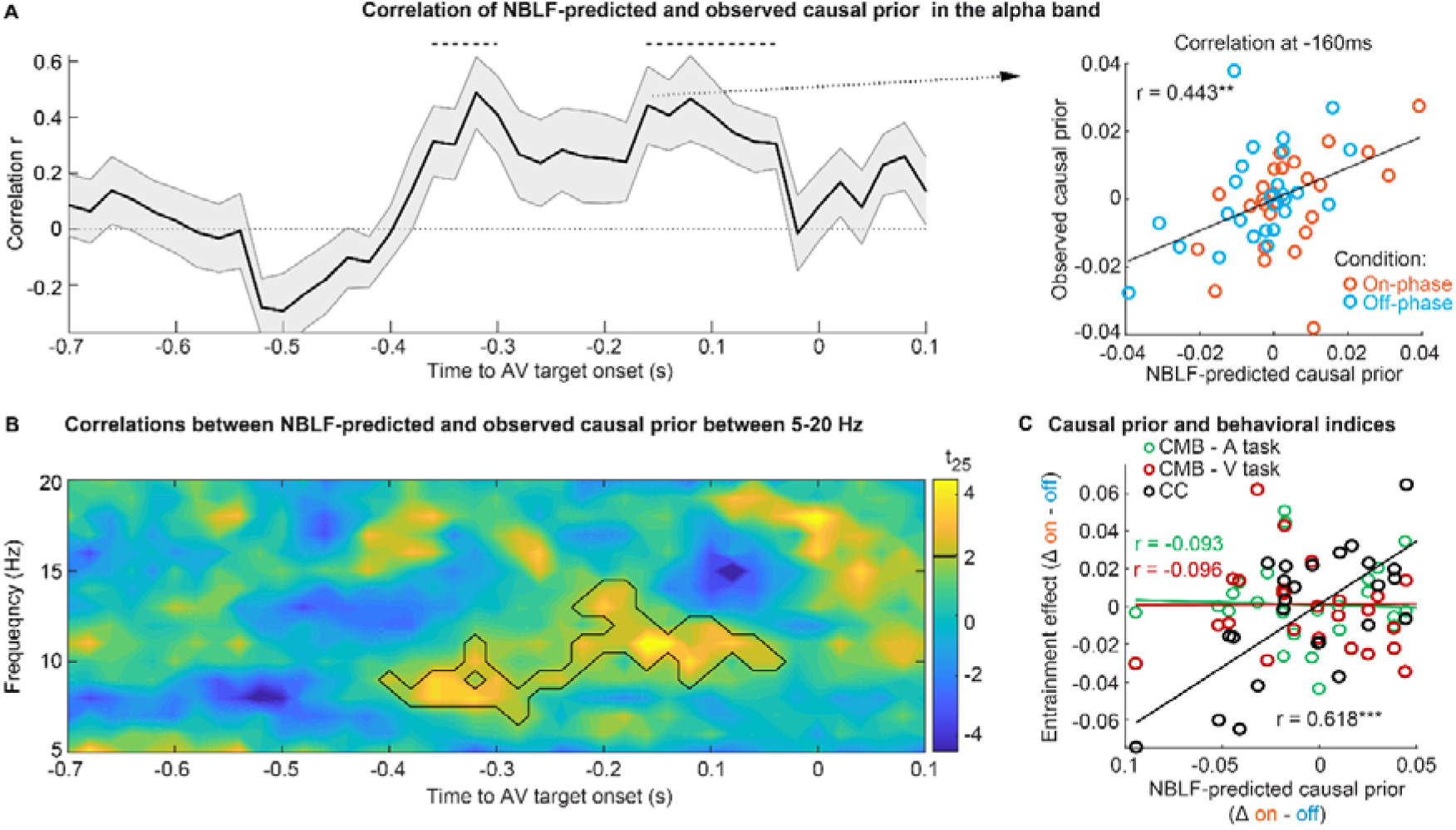
Individual neurobehavioural link functions predict the causal effect of alpha phase entrainment on audiovisual integration, particularly for causal judgments. **(A)** Correlations between the NBLF-predicted and empirically observed causal priors in the alpha band. The horizontal dotted lines indicate significant clusters from 360 to 300 ms and from 160 to 40 ms before AV target onset (p < 0.05, one-sided cluster-based randomisation t_25_ tests). The right panel shows the positive relationship between the NBLF-predicted and the observed multisensory causal prior at 160 ms before AV target onset. **(B)** Time-frequency t-map of correlations from 5-20 Hz. Significant clusters (p < 0.05; one-sided cluster-based randomisation t_25_ test) are marked by a solid line. The map shows a significant cluster between 400 to 40 ms before AV target onset, spanning a frequency range of 7-14 Hz. **(C)** The scatter plot at -160 ms shows a strong correlation (r = 0.618) between the predicted alpha (11 Hz) phase modulation of the multisensory causal prior and the main effects of alpha phase entrainment on common-cause judgements specifically in the causal judgement task (CC). *** = p < 0.001

Further analysis examined whether the modulation of the causal prior by the alpha phase differed between the causal judgements and the auditory and visual numerical report tasks. We correlated the behavioural indices from the three tasks with the alpha-phase-modulated causal prior from the significant alpha band cluster depicted in Fig. 6B. The causal prior modulation correlated highly with the causal judgements (r = 0.618, p < 0.001, BF_10_ = 40.722), but not with the CMBs computed from the auditory (r = -0.093, p = 0.652, BF_10_ = 0.167) or visual (r = -0.096, p = 0.639, BF_10_ = 0.168) numerical reports (Fig. 6C). These results suggest that the relationship between the causal prior and the alpha phase, when considering individual NBLFs, is driven by the causal judgement task. This task measures how the observers infer causality between multisensory signals. Taken together, these findings provide strong evidence for a causal effect of the individual alpha phase in multisensory integration.

### A psychophysical experiment showed individual behavioural effects of phase entrainment on multisensory integration at 5 and 10 Hz visual stimulation, but not for 15 Hz stimulation

Having demonstrated the individual effect of the alpha phase in multisensory integration, we wondered about the frequency specificity of this effect. We were also interested in investigating whether the 10 Hz entrainment effect could be replicated behaviourally, as well as exploring the possibility of describing the relationship between the phase of the entraining stimulus and audiovisual integration using a physical-behavioural link function (PBLF; see Methods). To this end, we conducted a follow-up multi-day psychophysical experiment in an independent sample of 29 observers. We adapted the EEG study setup to enable modelling of the individual PBLFs of the entraining visual stimulus in the theta (5 Hz), alpha (10 Hz), and beta (15 Hz) bands, in eight phase steps in relation to the AV target onset (Fig. 7A). As the PBLFs were derived directly from the entrainment conditions, there was no control condition.

**Figure 7.**
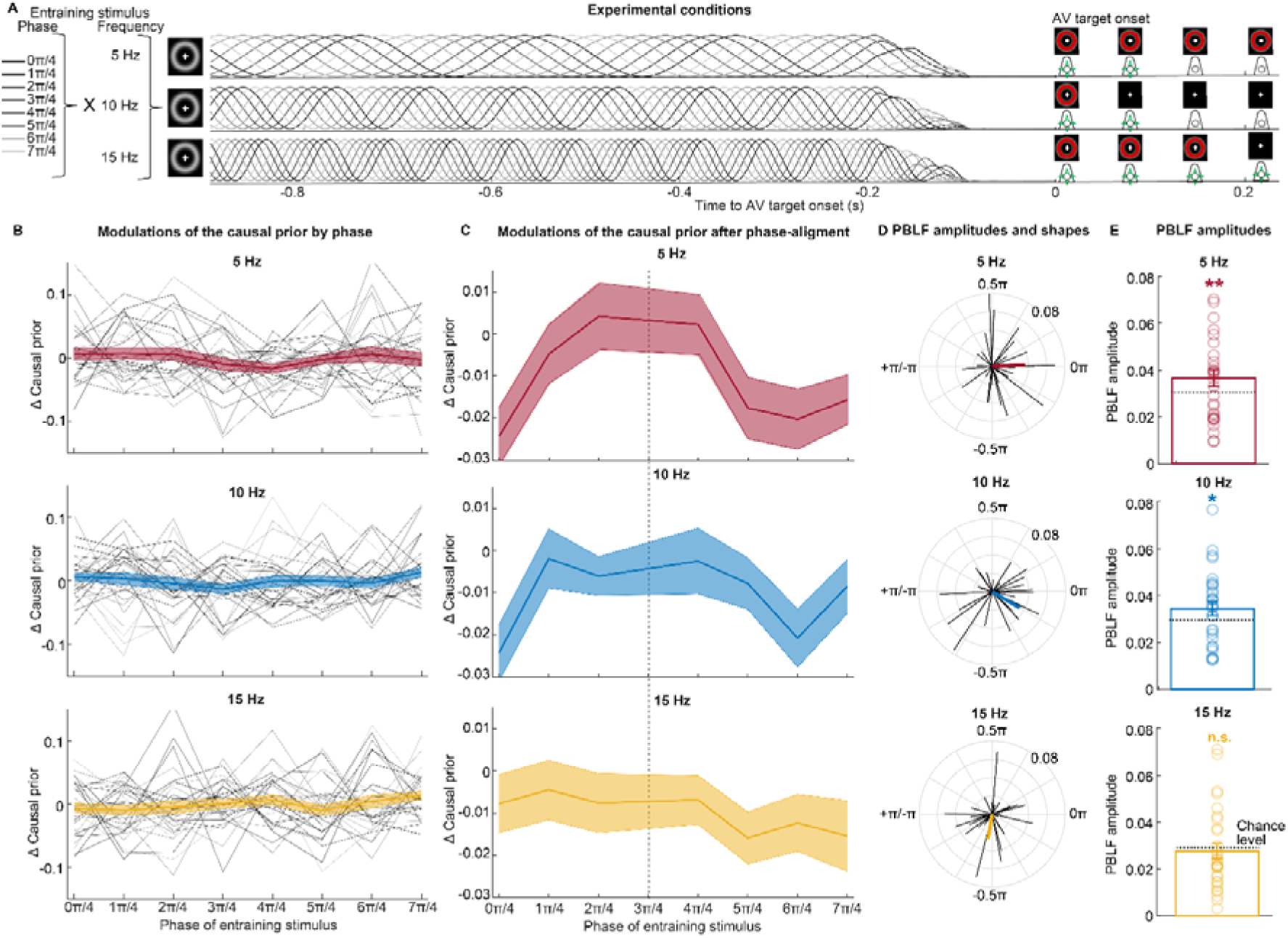
Findings of a psychophysical experiment show individual behavioural effects of phase entrainment on multisensory integration for 5 and 10 Hz visual stimulation, but not for 15 Hz. **(A)** Experimental setup for the 5, 10 and 15 Hz visual stimulation conditions. This behavioural study had a similar setup and tasks as the EEG study (Fig. 1), but no EEG was recorded. Rhythmic sinusoidal visual stimuli were instead presented at 5, 10 or 15 Hz at eight phase angles relative to the onset of the AV target stimuli. Entrainment effects were calculated separately for the three stimulation frequencies using a physical-behavioural link function (PBLF), which modelled the individual relationships between the phase of the entraining visual stimuli and the causal prior. **(B)** Modulations of the causal prior for the three stimulation frequencies (5 Hz, *top*; 10 Hz, *middle*; 15 Hz, *bottom*; the difference of the causal prior at each phase compared to the individual mean is shown) over eight phases of the entraining stimuli (x-axis). Across observers, no consistent effect of the phase of the entrainment stimulus on the causal prior was found for any frequency (the mean is depicted as bold traces, and the thin traces represent individual observers). **(C)** Average modulations of the causal prior, as shown in (B), after aligning the causal prior modulation by phase across participants (i.e. the individual maxima were shifted to 3π/4 (dotted vertical line; data point not plotted) prior to averaging). A sinusoidal modulation of the causal prior emerged for 5 and 10 Hz, but not for 15 Hz stimulation. **(D)** Amplitudes and shapes of the sinusoidal PBLF (individual, grey; average, coloured). For all three stimulation frequencies, the shapes were uniformly distributed around the circle, arguing against a generic PBLF (Raleigh tests: 5 Hz, z = 1.257, p = 0.287, BF_10_ = 0.024; 10 Hz, z = 1.232, p = 0.294, BF_10_ = 0.023; 15 Hz: z = 0.500, p = 0.611, BF_10_ = 0.011). **(E)** Amplitudes of the PBLF (individual and mean ± SEM). The dotted horizontal lines show the mean amplitude of the data for which the phase condition was randomised (n = 5000) prior to modelling the PBLFs. Significant PLBF amplitudes were found for the 5 Hz (*top*) and 10 Hz (*middle*) stimulation, but not for 15 Hz (*bottom*). This demonstrates the individual effect of entraining at 5 Hz and 10 Hz on multisensory integration, as reflected by the multisensory causal prior. Deviations from chance level are significant at p < 0.05 (*) and p = < 0.01 (**). Non-significant differences are indicated by “n.s.”.

When we fitted the BCI model to the behavioural data, we replicated the BCI modelling results from the EEG study (Table S4). The fitting procedure estimated the multisensory causal prior for each phase step and frequency (Fig. 7B). Repeated-measures ANOVAs revealed that the modulation of the causal prior by stimulus phase was non-significant across observers, as the causal priors were equivalent across the eight phase steps of the entraining stimuli at all three frequencies (see supplementary results). Yet, when the causal prior was aligned by individual phases of the observers^30^, a sinusoidal pattern emerged for 5 and 10 Hz, but not for 15 Hz (Fig. 7C). Interestingly, modelling the individual causal priors (Fig. 7B) over the eight phase steps as individual sinusoidal PBLFs revealed that the shapes of the PBLFs were uniformly distributed around the circle (Fig. 7D and Fig. S8), which provides clear evidence against a generic PBLF. However, the PBLF amplitudes were significantly larger than expected by chance level for the 5 Hz (t_28_ = 2.783, p = 0.006, Cohen’s d = 0.515, BF_10_ = 4.644) and 10 Hz (t_28_ = 2.176, p = 0.020, Cohen’s d = 0.407, BF_10_ = 1.549) entrainment conditions (Fig. 7E). This was not true for the 15 Hz entrainment condition (t_28_ = -0.540, p = 0.704, Cohen’s d = 0.099, BF_10_ = 0.225). Furthermore, the PBLF amplitudes were larger at 5 Hz than at 15 Hz (t_28_ = 1.656, p = 0.041, Cohen’s d = 0.307, BF_10_ = 0.664) and 10 Hz than at 15 Hz (t_28_ = 1.626, p = 0.045, Cohen’s d = 0.302, BF_10_ = 0.637). PBLF amplitudes were equivalent for 5 Hz and 10 Hz (t_28_ = -0.54894, p = 0.547, Cohen’s d = 0.102, BF_10_ = 0.227). Since PLBFs were computed from the entrainment conditions and not from an independent control condition (as in the EEG study), no correlations were computed between the PBLF-predicted and the observed entrainment effects on behavioural multisensory integration indices, because this would be circular^80^. Overall, the follow-up psychophysical experiment showed that phase entrainment has a direct behavioural effect on multisensory integration in the theta (5 Hz) and alpha (10 Hz) bands when individual phase-behaviour relationships were modelled with PBLFs. Yet, this effect was not observed for the beta (15 Hz) band, indicating an upper frequency boundary for individual phase-entrainment effects on audiovisual perception.

## Discussion

In two studies, we examined the causal role of the individual alpha phase in multisensory integration. In a multi-day computational EEG study, we entrained the alpha phase using rhythmic visual stimulation and modelled audiovisual perception with an established multisensory paradigm^19,51^. In a follow-up psychophysical experiment, we investigated the frequency specificity of the individual phase entrainment effects observed in the EEG study. As part of this work, we developed the NBLF as a novel analytical approach for probing individual causal relationships between the alpha phase and perception. Specifically, NBLFs enabled us to test the direct causal path from the entrained alpha phase to perception, thereby bypassing limitations of previous studies which may have been affected by nonspecific neural mechanisms.

In line with our previous experiments^19,51^, as well as reports on the sound-induced flash illusion^57^ and other multisensory signal combinations^58,60,81–84^, our behavioural data revealed key profiles of multisensory integration effects in numerical reports and causal judgements. The profiles were well fitted by the BCI model^52,53^.The parameters of the fitted BCI model, including the multisensory causal prior, did not differ significantly between the on-phase and off-phase entrainment conditions, with moderate to strong evidence for their equivalence. The absence of entrainment effects on perception suggests that there is no generic relationship between the alpha phase and audiovisual integration. This is consistent with the conflicting results of previous sensory entrainment studies^46^, in which some studies demonstrated phase entrainment effects on visual detection and discrimination^37–39,42–44^, while others did not^40,41,45^. Thus, at the group level, the behavioural data revealed no differential effects of on-phase and off-phase 10 Hz visual stimulation on perception. To better understand the behavioural results and to test for a causal effect of the alpha phase manipulation on perception, it was necessary to demonstrate that the 10 Hz visual stimulation entrained endogenous alpha oscillations. We thoroughly examined the effects of the 10 Hz stimulation on different EEG markers, including power, ITPC, individual and instantaneous alpha frequency, and 10 Hz phase. In line with previous studies^39,41,47,54,70^, we found that the 10 Hz visual stimulation reliably entrained alpha oscillations in the early visual cortex. Furthermore, our analysis revealed clear evidence that the observed 10 Hz response pattern in the EEG reflects an entrainment of endogenous alpha oscillations rather than a superposition of event-related potentials^64–68,73^. Notably, alpha oscillations in the on-phase and off-phase conditions were precisely entrained with a constant phase angle difference of 180° between conditions, showing that our experimental manipulation precisely altered the phase of alpha oscillations.

Having established that the 10 Hz visual stimulation entrained the phase of endogenous alpha oscillations, we computed the NBLF for each observer separately from data of the control condition. The significant NBLF amplitude observed from 340 to 180 ms before AV target onset in the non-entrainment control condition was consistent with the findings of our previous study^19^. Notably, the shape of the NBLFs varied considerably between observers, providing clear evidence of individual relationships between the alpha phase and multisensory integration. Our findings are consistent with the results of previous studies that identified interindividual differences in phase-behaviour relationships in auditory^31,32^, visual^14,33,34^, somatosensory^35^ and audiovisual^19^ perception. However, while the data from the control condition revealed a correlation between the individual alpha phase and multisensory integration, they did by not prove causality. Therefore, it was necessary to demonstrate that the individual entrainment effects on perception, as predicted by the NBLFs, are related to the observed entrainment effects. This was the case. The predicted and the observed entrainment effects on audiovisual perception showed a positive relationship from 360 ms to 40 ms before the AV target onset, demonstrating an individual causal effect of the alpha phase in multisensory perception.

Interestingly, the causal effects of the alpha phase mediated by the NBLF were particularly evident in the causal judgement task, but not in the numerical report tasks. This suggests that specific task demands are necessary to establish this relationship. The causal judgement task required top-down bisensory attention, which has been shown to facilitate audiovisual integration^85,86^ and alter network coupling in the alpha band^28,87^. In the current study, the NBLF was primarily associated with alpha phase coupling between the early auditory and visual cortices, aligning with these reports. As observers are explicitly asked to judge multisensory integration in the causal judgement task, our findings imply that top-down attention modulates multisensory processing via modulation of the alpha phase coupling between early auditory and visual cortices. This coupling presumably plays a role in forming individual alpha phase-perception relationships. Importantly, our findings also suggest that the alpha phase not only reflects periodic alterations in excitability that modulate local information processing^5,7,11,13^, but also plays a role in regulating information flow between different cortical regions^88,89^. For example, a recent optogenetic study revealed that the alpha cycle in the visual cortex alternates between a depolarised state, employed for internal processing, and an excitable state, employed for detecting external inputs from other cortical regions^8^. Together, these findings establish an individual causal role of the alpha phase in multisensory integration. This relationship depends on task demands and was associated with alpha phase coupling between early audio-visual areas.

To further investigate whether the individual alpha phase-perception relationships could be replicated at the behavioural level and were specific to 10 Hz visual stimulation, we conducted a follow-up psychophysical study. This study revealed individual phase-perception relationships for 5 and 10 Hz visual stimulation, but not at 15 Hz. These results are consistent with previously reported entrainment effects on unisensory visual discrimination at 5 Hz and 10 Hz, but not at 15 Hz^38^. However, it is possible that the 5 Hz visual stimulation entrained alpha oscillations at the second harmonic (i.e. 10 Hz), and that this entrainment mediated the effect of 5 Hz visual stimulation on multisensory perception in our study. In contrast, rhythmic stimulation at 15 Hz may not have reliably entrained endogenous oscillations relevant for perception^70^. Future electrophysiological studies could investigate whether 10 Hz neural harmonics in response to 5 Hz visual stimulation have an individual effect on multisensory integration or whether the observed effect is explained by entrainment of the endogenous 5 Hz phase by itself. Notably, theta oscillations around 5 Hz have been suggested to modulate attentional sampling^29^, which influences the multisensory causal prior^90^. Therefore, the 5 Hz visual stimulation in the psychophysical study may have influenced audiovisual integration by modulating attentional sampling, a hypothesis that could be targeted in further studies using the NBLF approach. Taken together, our psychophysical study revealed direct individual relationships between the phase of entraining 5 and 10 Hz visual stimulation on multisensory perception. The absence of such effects for 15 Hz stimulation indicates an upper frequency boundary for phase entrainment effects on audiovisual perception.

As a limitation of this study, it is important to emphasise that other factors, which cannot be directly measured by EEG, such as the precise configuration of the alpha network, conduction delays between regions, and cortical folding^11,33,91^, may also have contributed to the interindividual differences in alpha phase effects on perception. For instance, variations in alpha network configurations^91^ could explain why alpha oscillations are detected differently in scalp-recorded EEG data. If this were the case, one would expect to find interindividual differences in phase-perception relationships even in the presence of generic NBLFs. However, we consider this unlikely, given that individual phase-perception relationships were identified for both the alpha phase in the EEG study and the phase of the entraining visual stimuli in the psychophysical study. This suggests that the individual relationships between the alpha phase and perception did not only arise from limitations of the EEG measurement. Nevertheless, differences between individuals in conduction delays and cortical folding, which influence the timing and processing of stimuli, may play a role in forming these individual relationships. Future intracranial studies could shed new light on this question^92,93^. Furthermore, high-field magnetic resonance tomography and diffusion tensor imaging studies could reveal how anatomical factors contribute to individual alpha phase-perception relationships. In summary, while our study showed that the alpha amplitude and phase coupling between early audio-visual cortices contribute to the individual phase-perception relationships, other factors, such as conduction delays and anatomical differences may also account for the existence of individual NBLFs and PLBFs.

Our results have important theoretical, methodological and clinical implications. They suggest that multisensory integration operates periodically, with the waxing and waning of alpha oscillations providing temporal windows for the up- and downregulation of the multisensory information flow within cortical networks. Such perceptual cycles have been suggested previously for unisensory visual perception^10–13^. To account for the interindividual variability in the relationship between alpha phase and perception, future MEG/EEG studies should model NBLFs at the individual level. Without explicitly considering individual NBLFs, existing causal phase effects may be overlooked when averaging data across observers. Our findings also have implications for neurotechnologies and personalised medicine. For instance, studies employing non-invasive closed-loop brain stimulation or brain-computer interfaces^94–96^ to utilise or target the alpha phase should consider individual phase-perception relationships. Taking these relationships into account may also lead to a better outcomes in clinical research examining oscillatory phase as a personalised disease biomarker^97^. Overall, our study highlights the importance of modelling individual brain-perception relationships to advance personalised clinical and cognitive neuroscience research^97,98^.

## Methods

### Participants

Twenty-six healthy volunteers participated in the EEG study after providing written informed consent. An additional sample of 30 healthy volunteers participated in the follow-up psychophysical study. These participants numbers were based on prior statistical power calculations, for which we aimed to statistically secure medium effects size (Cohen’s d ∼ 0.6) estimated from our previous study^19^. For the psychophysical experiment, we estimated the sample size from the results of the current EEG study. Participants were recruited at local universities. Inclusion criteria: Age 20-65 years; normal or corrected to normal vision and hearing. Exclusion criteria: Psychiatric, neurological or psychosomatic disorders; cardiovascular disorders; and diabetes. One participant was excluded from the psychophysical study because of their accuracy in visual and auditory numerical reports was low (< 3 interquartile ranges below the median). A total of 26 participants (EEG study: mean age 25.5 years, range 18-37 years; 16 females) and 29 participants (psychophysical study: mean age 22.0 years; range 18-41 years; 21 females) were included in the final data analysis. The studies were approved by the local human research review boards of the University of Tübingen (EEG study; approval number 200/2017BO2) and the Friedrich-Alexander-Universität Erlangen-Nürnberg (psychophysical study; approval number 20-525_1-B).

### Stimuli

The setups of the two studies differed slightly because the follow-up study presented different entrainment frequencies and different latencies between entrainment stimuli offsets and target stimuli onset. Importantly, this adjustment did not markedly affect the behavioural outcomes of the critical 10 Hz entrainment effects. Stimuli were controlled using Psychtoolbox (PTB^99,100^; www.psychtoolbox.org), version 3.09 for the EEG study and version 3.0.19 for the psychophysical study, running under MATLAB (MathWorks), version R2016a for the EEG study and version R2020a for the psychophysical study. In the EEG study, PTB sent trigger pulses to the EEG recording system. Auditory stimuli were presented at ∼75 dB SPL via two loudspeakers (EEG study: Logitech Z130; psychophysical study: Speedlink Twoxo), which were positioned on each side of the monitor (EEG study) or above the monitor (psychophysical study). Visual stimuli were presented on an LCD screen with a 60 Hz refresh rate (EEG study: EIZO FlexScan S2202W) or with a 100 Hz refresh rate (psychophysical study: ASUS ROG Swift PG278Q). Button presses were recorded using either a standard keyboard (EEG study) or a four-button response pad (The Black Box ToolKit Ltd.). Participants sat in front of the monitor and loudspeakers at distances of 85 cm (EEG study) and 61 cm (psychophysical study). Both studies were conducted in a sound-attenuated room. For the EEG study, the room was also electrically shielded.

In both studies, the entraining visual stimulus (Fig. 1) was a white circle presented in the centre of a black screen. The circle’s maximum greyscale value (i.e. white) was at a radius of 5°. The circle’s smoothed inner and outer borders were defined using a greyscale value that followed a Gaussian distribution with an STD of 1.25°. In the EEG study, to entrain 10 Hz oscillations, the circle’s contrast was sinusoidally modulated between 0% and 100% contrast at 10 Hz (on-phase and off-phase conditions), or kept constant at 50% contrast (control condition). In the psychophysical study, the circle’s contrast was modulated sinusoidally at 5, 10, or 15 Hz. The phase of the sinusoidal modulation was manipulated relative to the onset of AV target stimuli at two opposite mean phase angles (EEG study: 0π vs. 1π, i.e. peak vs. trough of sinusoid at -200 ms) or eight equidistant phase angles (psychophysical study: step size π/4 from 0π to 7π/4). The onset of the entraining circles was jittered between -933 and -533 ms (EEG study) or between -1000 ms and -700 ms (psychophysical study) before the AV target stimulus onset. The offset of the circle modulation was jittered between -166 ms and -50 ms (EEG study) or between -200 ms and - 100 ms (psychophysical study) before the AV target stimulus onset. Thus, the duration of the sinusoidal modulation of the circle was between 500 ms and 800 ms (EEG study) or between 600 ms and 800 ms (psychophysical study). In the EEG study, after the sinusoidal modulation offset, the circle was presented at 50% contrast until -33 ms before the AV target stimulus onset. Then, a black screen was presented for 33 ms duration until the AV target stimulus onset. In the psychophysical study, a black screen was presented after the offset of the sinusoidal modulation of the circle (i.e. jittered between 100 to 200 ms). The timing of the stimuli was verified using photodiode and microphone measurements.

In adaptation of a sound-induced-flash-illusion paradigm^55–57^, the visual stimuli of the AV target stimuli consisted of a red circle presented in the center of the black screen (100% contrast with RGB values (200, 0, 0; Fig. 1A) for 16.7 ms in the EEG study and for 20 ms in the psychophysical study. The circle’s maximum red value (i.e. RGB = 200) was at a radius of 5°. The circle’s smoothed inner and outer borders were defined using a red value that followed a Gaussian distribution with an STD of 1°. The auditory beep was a 2000 Hz pure tone with a duration of 16.7 ms (EEG study) or 20 ms (psychophysical study). Its amplitude was modulated by a Gaussian envelope centred at the middle of the beep to create an on/off amplitude ramp. One to four visual flashes and one to four auditory beeps were presented sequentially at a fixed SOA of 66.7 ms (EEG study) or 70 ms (psychophysical study). Within an AV flash-beep sequence, each flash and/or beep was presented in a fixed temporal slot starting at 0, 66.7, 133.3 or 200 ms (EEG study) or 0, 70, 140 or 210 ms (psychophysical study). The temporal slots were filled sequentially. If the same number of flashes and beeps were presented on a particular trial, the beeps and flashes were presented in synchrony. On trials with different numbers of beeps and flashes, the additional beeps (or flashes) were presented in subsequent fixed time slots.

### Experimental setup

Participants were instructed to report either the number of flashes (i.e. visual numerical report task) or the number of beeps (i.e. auditory numerical report task) in different blocks, while ignoring inputs in the task-irrelevant modality. In a third task, which explicitly required attention to be given to both sensory modalities, participants had to judge whether both stimuli (i.e. auditory and visual input) derived from one common cause or from independent causes (i.e. causal judgement task). The EEG study used a 3×4×4×3 factorial design that manipulated the alpha entrainment condition (control condition, on-phase, or off-phase stimulation), the number of visual flashes (1 to 4), the number of auditory beeps (1 to 4), and the task (auditory numerical report, visual numerical report, causal judgement). This yielded a total of 144 AV conditions (Fig. 1B). For the crossmodal bias analyses, the trials were reorganized based on their absolute numeric disparity (|#A - #V| ∈ {0,1,2,3}; Fig. 2). The psychophysical study used a 3×8×4×4×3 factorial design that manipulated entrainment frequency (5 Hz, 10 Hz, 15 Hz), entrainment phase (8 phase steps), the number of visual flashes (1 to 4), the number of auditory beeps (1 to 4), and the task (auditory numerical report, visual numerical report, causal judgement). This yielded a total of 1,152 AV conditions. Both studies included unisensory conditions and the psychophysical study additionally included AV conditions without an entraining visual stimulus in a separate block as control conditions. A response screen was presented between 50 ms to 200 ms (EEG study) or 150 ms to 250 ms (psychophysical study) after the offset of the last flash/beep stimulus. The response screen presented textual instructions to report the perceived number of flashes or beeps, or to provide a causal judgement. Participants responded as accurately as possible by pushing a respective response button within 2.5 s. The order of buttons (ascending numbers left to right or vice versa) was counterbalanced across runs to decorrelate motor responses from numerical reports and causal judgements. After a participant’s response, the onset of the entraining visual stimuli in the next trial started after a jittered fixation interval of 0.57–1.7 s (EEG study) or 0.77-3.77 s (psychophysical study).

In each AV block of the EEG study, participants were presented with 170 trials in a pseudo-randomised order. Each of the 16 AV target stimuli (i.e. 1-4 flashes × 1-4 beeps) in the control condition was presented 4 times and each of the 16 AV target stimuli in the on-phase and off-phase entrainment conditions was presented 3 times. Additionally, 10 ‘null trials’ were interspersed within each block. In these trials, neither the static circle in the control condition nor the entraining stimuli in the on-phase or off-phase conditions were followed by an AV target stimulus. Participants completed 6 blocks of each of the auditory numerical report, visual numerical report, and causal judgement tasks (i.e. 18 blocks in total). These 18 blocks were presented in a counterbalanced order and were followed by two unisensory blocks with visual or auditory target stimuli only. Data from the unisensory blocks were used for the behavioural control analyses (Fig. S2). In the unisensory blocks participants were presented with 141 trials from 12 unisensory conditions (i.e. 4 signal numbers x 3 alpha entrainment conditions), each of which was presented 11 times per block. In addition, 3 types of ‘null trials’ were presented 3 times per block. Data were collected in two EEG-recording sessions on different days. Participants completed a total of 20 blocks, for a total of 3,342 trials. On average, the duration of data collection in each EEG session was 92 minutes.

In the psychophysical experiment, participants were presented with 128 trials in blocks with AV target stimuli after the entraining V stimulus, with 80 trials in runs with AV target stimuli *without* entraining V stimulus, and with 32 trials in blocks with unisensory conditions. Stimuli from different conditions were presented in pseudorandom order. For AV blocks involving an entraining visual stimulus, participants completed 6 runs each of auditory report and visual report, and 6 runs of causal judgements. As in the EEG study, the 18 AV blocks were presented in a counterbalanced order, followed by two unisensory blocks containing visual or auditory target stimuli only. The 1,152 AV conditions were presented twice across the 18 AV blocks. After the AV blocks, participants completed 3 additional AV blocks without any entraining visual stimulus with auditory numerical report, visual numerical report, and causal judgements. Additionally, participants completed 2 unisensory runs with visual or auditory stimuli only and without any entraining visual stimulus. In AV blocks without an entraining visual stimulus, participants were presented with 80 trials from 16 AV conditions (i.e. 1-4 flashes x 1-4 beeps) each 5 times. In unisensory runs, participants were presented with 32 trials from 4 unisensory conditions (i.e. 1-4 flashes or beeps without an entraining V stimulus), each 4 times. Over two experimental sessions on different days, participants completed 23 blocks, for a total of 2,608 trials (except for one participant with only 1,705 trials due to technical issues). The duration of data collection for each psychophysical study session was about 108 minutes. In both studies, participants began with 46 practice trials.

### Behaviour - GLM-based analyses in the EEG study

In the EEG study, GLM-based analyses were used to examine participants’ auditory and visual reports, as well as their causal judgements regarding AV target stimuli. For the auditory and visual numerical reports, the crossmodal bias (CMB) was computed based on the numerical reports of AV incongruent stimuli (i.e. trials in which a different number of visual and auditory stimuli were presented), in order to characterise how participants weigh audiovisual signals during multisensory perception^51,58,59^. The CMB quantifies the relative influence of the auditory (n_A_) and the visual (n_V_) target stimuli on observers’ auditory and visual numerical reports (r_A/V_):

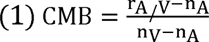

To account for response biases, n_A_ and n_V_ were adjusted using a linear regression approach in a participant-specific manner^58^. The participants’ responses were regressed against the number of AV stimuli in all congruent trials. The true n_A_ and n_V_ in the CMB equations were replaced with the n_A_ and n_V_ values predicted by the fitted linear regression model. The CMB ranged between zero (i.e. pure auditory influence) and one (i.e. pure visual influence).

Individual CMBs were analysed using repeated-measures ANOVAs with the following within-participant factors (Tab. S1): absolute numeric disparity (1, 2, 3); task relevance (auditory vs. visual report); and alpha entrainment condition (control, on-phase, off-phase). For causal judgements, the proportion of common-cause responses was computed, and repeated-measures ANOVAs were performed with the within-participant factors of absolute numeric disparity (1, 2, 3) and alpha entrainment condition (control, on-phase, off-phase). To assess differences between the on-phase and off-phase alpha entrainment conditions, these ANOVAs were repeated with the factor entrainment condition with only two levels (on-vs. off-phase). Complementary Bayesian repeated-measures ANOVAs were used to provide evidence that there was no difference between the on-phase and off-phase alpha entrainment conditions. Bayesian ANOVAs allow one to compute inclusion Bayes factors (BF_Incl_) for each experimental factor. These quantify the average extent to which including the factor in the ANOVA models improves their fit, despite the increased complexity of these models^101,102^. All classical and Bayesian ANOVAs were computed using JASP 0.17.2.1^102^.

### Behaviour - Causal inference model

To investigate whether participants adopted a BCI or simpler heuristic decision strategies^19,52,103^, five competing models were fitted to participants’ visual and auditory numerical reports, as well as their causal judgements. In the EEG study, these models were fitted to the behavioural data of all three tasks, separately for each of the three experimental conditions (i.e. control, on-phase, off-phase) to compare the competing models and model parameters between conditions. To model the modulation of the causal prior by alpha phase with NBLFs in the non-entrainment control condition, and to make predictions for the two entrainment conditions, the BCI model was fitted to all three experimental conditions in order to determine the model that best fit the data overall.

In the psychophysical experiment, the competing models were fitted across all AV conditions at an entrainment frequency (5, 10 or 15 Hz). Specifically, one causal prior parameter was fitted for each of the 8 distinct phase modulations. Physical-behavioural link functions (PBLFs) were then fitted to causal prior parameters of the fitted BCI model. In both studies, Bayesian model comparison was used to determine the model that best explains the observers’ behavioural data. Details on the BCI model, the fitting to behavioural judgements and Bayesian model comparisons have been provided elsewhere^19,52,103^, and are summarized in the supplemental methods.

A Bayesian optimization algorithm, as implemented in the BADS toolbox^104^, was used to obtain maximum likelihood estimates for the models’ parameters. In the EEG study, 5 parameters (pCommon, μ_P_, σ_P_, σ_A_, σ_V_) were fitted for the BCI model. In the psychophysical study, 4 basic parameters (μ_P_, σ_P_, σ_A_, σ_V_) plus 8 parameters of the causal prior were fitted for each of the 8 entrainment phases (pCommon_1_, pCommon_2_, …, pCommon_8_). The BADS optimization algorithm was initialised 20 times for the EEG study and 25 times for the psychophysical study. The results (i.e. model comparisons and parameters) are reported for the models with the highest log likelihood across these initialisations (Tab. S2).

To compare the BCI model’s fitted parameters between the three experimental conditions in the EEG study, randomisation tests (i.e. under exchange of experimental conditions) were used with t-values as test statistics (n = 5,000 randomisations). To assess the evidence in favour of the null hypothesis (i.e. no difference between conditions), the randomisation tests were complemented by Bayesian t-tests using the BayesFactor toolbox^105^ to quantify evidence in favour of a condition difference (H1) relative to the null hypothesis of no difference (H0).

### EEG - Data acquisition and preprocessing

EEG signals were recorded from 64 active electrodes positioned in an extended 10–20 montage using electrode caps (actiCap, Brain Products, Gilching, Germany) and two 32 channel DC amplifiers (BrainAmp, Brain Products). Electrodes were referenced to FCz using AFz as ground during recording. Signals were digitized at 500 Hz with a high-pass filter of 0.1 Hz. Electrode impedances were kept below 25 kOhm. Preprocessing of EEG data was performed using Brainstorm^106^ running on Matlab R2021a. Eye blinks were automatically detected using data from the FP1 electrode (i.e. a blink was detected if the band-pass (1.5–15 Hz) filtered EEG signal exceeded two times the STD; the minimum duration between two consecutive blinks was 800 ms). Signal-space projectors (SSPs) were created from band-pass filtered (1.5–15 Hz) 400 ms segments centred on detected blinks. The first spatial component of the SSPs was then used to correct blink artifacts in continuous EEG data. Further, all data were visually inspected for artifacts from blinks (i.e. residual blink artifacts after correction using SSPs), saccades, motion, electrode drifts or jumps. On average 8.6% + 2.3% SEM of AV trials were affected. However, because our main analyses focused on electrodes over the occipital cortex (Fig. 3A) which were largely unaffected by artifacts, we used all trials in our EEG analyses. A control analysis showed that ERPs and ITPC (Fig. 4A) were highly similar if we excluded affected epochs.

For the ERP data analysis, the EEG was band-pass filtered (2–45 Hz), re-referenced to the average of left and right mastoid electrodes (TP9 and TP10), and downsampled to 150 Hz. ERPs were analysed from 1 s before AV target stimulus onset up to 0.5 s after stimulus onset and all epochs were baseline corrected using a prestimulus baseline from -1.4 to -1 s (i.e. including time points before the onset of the entraining V stimulus). To analyse an occipital oscillator, the EEG data from six electrodes located over the visual cortex (i.e. O1, O2, Oz, PO3, POz, PO4) were averaged. For the time-frequency analyses, including modelling of NBLFs and their prediction of causal prior modulations, continuous EEG data were band-pass filtered from 0.25 to 85 Hz with a notch filter at 50 Hz and epoched from -2.4 to 1.5 s relative to AV target onset. The spectral power and phase of single-trial EEG data were extracted using complex Morlet wavelets (as implemented in Brainstorm). This was done from -1.4 s to +0.5 s in 1 Hz steps from 5 to 24 Hz and in 2 Hz steps from 26 to 40 Hz, with the number of wavelet cycles increasing linearly from 4 to 10 across frequencies^107^ and a full width at half maximum of 0.5. Finally, all time-frequency representations were downsampled to 50 Hz.

### EEG - Analysis of entrainment of occipital oscillator

ERPs in response to the 10 Hz visual stimuli were analysed at both the scalp and source level (Fig. 3). Source localization was performed using a minimum norm estimation with sLoreta and an OpenMEEG BEM head model. A noise covariance matrix was computed from -1.4 to -0.9 s before the AV target stimulus onset, as implemented in Brainstorm. To plot source-level data in Fig. 3B, data were extracted from early visual areas (V1/V2; Brodmann’s atlas in Brainstorm) and bandpass-filtered between 8 to 12 Hz. To further investigate the entrainment, inter-trial phase clustering (ITPC) was computed at 5 to 40 Hz for each time-frequency point, representing the absolute length of the mean phase angle vector^107^. The mean phase angle vector was computed across trials separately for each individual. Next, ITPC was baseline-corrected by subtracting the average ITPC from a prestimulus baseline time window between -1.4 to -1.3 s before the AV target stimulus onset, for each frequency. The baseline-corrected ITPC was then tested against zero at the group level using a nonparametric randomisation test (5,000 randomisations) in which the sign of the normalised ITPC was flipped^108^. One-sided one-sample t-tests were computed as the test statistic, corrected for multiple comparisons across time-frequency points using a cluster-based correction for p-values^109^. The sum of t-values across a cluster was used as the cluster-level statistic and an auxiliary cluster-defining threshold of t = 5 was employed. To plot the evolution of phase angles’ entrainment, the distribution of phase angles at 10 Hz was computed between 0 and 2π for each time point in each individual, and then it was averaged across participants. Additionally, ITPC and phase distributions were computed using source-localised, time-frequency transformed data from the left and right V1 and V2 (Fig. S4).

Next, it was tested whether the 10 Hz stimulation entrained ongoing endogenous oscillations. These analyses investigated 5 to 20 Hz power, IAF, and phase angle evolution (Fig. 4). Power at each frequency was computed as the mean square length of the complex vector^107^ and was baseline-corrected as relative percent change in power using an interval of - 1.4 to -1.3 s before the AV target onset. The relative power was then tested against zero at the group level using nonparametric randomisation tests (5,000 randomisations with sign flipping). One-sided one-sample t-tests were used as the test statistic, with a cluster-based correction applied using a cluster-defining threshold of *t* = 2. Depending on the IAF of the participants, the power and ITPC were investigated by computing the individual peaks in the alpha band (7-13 Hz)^110^ across 6 electrodes over the visual cortex (Fig. 3A). This was achieved using two two-minute resting-state EEG data recordings with the eyes closed, obtained in between blocks of the main study. Power, ITPC and instantaneous alpha frequency (InstAF) were analysed in two groups, one with an IAF above 10 Hz (n = 16; 10.821 ± 0.786 Hz; mean ± STD IAF) and one with an IAF below 10 Hz (n = 10; 9.294 ± 0.359 Hz). As previously suggested^111^, the InstAF was calculated using band-pass filtered time-domain ERP epochs with the band-pass filter centred on the IAF (± 1 Hz). Filtered data were then Hilbert transformed to obtain the derivative of the unwrapped phase angle time series. To correct for phase slips due to noise, the median of ten running median filters was used, with a window size ranging from 20 to 400 ms, to obtain the final estimate of InstAF. The InstAF was averaged over occipital channels, and the entrainment versus control condition contrast was compared between the two groups with IAF above or below 10 Hz. A nonparametric randomisation test (i.e. 5,000 random group assignments) with a two-sided two-sample t-tests was used as test statistic. The outcome was cluster-based corrected with a cluster-defining threshold of t = 2. To investigate the time course of 10 Hz phase angles in ‘null trials’, the mean resultant length of the alpha phase and the mean phase angles in the on-phase and off-phase entrainment conditions were computed, along with their circular difference using the CircStat toolbox^112^.

### Modelling of neurobehavioural link functions and predictions of causal alpha phase effects

NBLFs were estimated between the alpha phase and the multisensory causal prior of the BCI model as a sinusoidal modulation of the constant causal prior pCommon_const_, which was fixed during the initial fitting procedure (supplemental methods and Tab. S2). The NBLF approach enabled the causal prior to be modelled as a trial-wise continuous quantity that varies with the trial-wise alpha phase, for a given time point and frequency (i.e. pCommon_i,t,f_). Continuous modelling of the causal prior enabled fine-grained predictions of the NBLFs, as opposed to the coarse and discrete predictions used in conventional sort-and-bin approaches^19^. Importantly, regression-based methods such as NBLFs are robust to violations of sinusoidality^30^. As alpha phase-behaviour relationships are sinusoidal, the mapping between phase and behaviour must be explicitly modelled in order to test whether an entrained alpha phase affects the causal prior: Only such an explicit mapping of NBLFs can accurately predict how a manipulated phase will lead to a specific high or low causal prior according to a ‘preferred’ or ‘non-preferred’ phase. In stimulation studies that entrain phase, failing to model individual NBLFs will likely result in false-negative null effects at the group level (see supplemental results and Fig. S8 for a more detailed discussion).

The NBLF between phase (computed from time-frequency transformations) at visual electrodes (i.e. O1, O2, Oz, PO3, POz, PO4) and the multisensory causal prior was fitted using data from the non-entrainment control condition only. For each time point (t) and frequency (f) in time-frequency space from -0.7 to 0.1 s before and after the AV target stimulus onset, the causal prior was fitted, in addition to the constant causal prior parameter (pCommon_const_), as a function of phase (C_i,t,f_) across all trials (i) using a sinusoidal function:

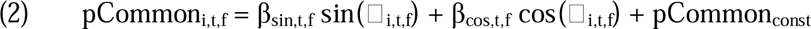

Thus, the NBLF amplitude and shape were determined by the two parameters β_sin,t,f_ and β_cos,t,f_. Specifically, the predicted distributions were computed for the auditory and visual numeric estimates (i.e. 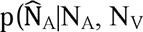) and 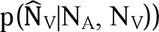 and for the posterior probability of a common cause (i.e. 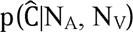). This was done for each trial, given the fixed parameters of the BCI model (pCommon_const_, μ_P_, σ_P_, σ_A_, σ_V_; as estimated in the initial fitting procedure across all conditions) and the variable causal prior (pCommon_i,t,f_). These distributions were generated by simulating x_A_ and x_V_ (i.e. continuous variables sampled from Gaussian distributions) 500 times for each trial and inferring 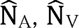, and 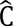 from the BCI model equations (see equations (1)-(9) in supplemental methods). As described for the main model, auditory and visual numerical reports, as well as causal judgements, were linked to the predicted distributions. From the predicted multinomial distributions (i.e. one for each trial), the log likelihoods were summed across all trials (i.e. 1152 trials) in the non-entrainment control condition.

A Bayesian optimization algorithm, implemented in the BADS toolbox^104^, was used to obtain maximum likelihood estimates of the NBLF parameters β_sin,t,f_ and β_cos,t,f_. The effect of oscillatory phase on the causal prior variable at a given point in time-frequency space (i.e. equation (2)) was estimated to optimize the prediction of auditory and visual numerical reports as well as causal judgements across all trials. The parameters β_sin,t,f_ and β_cos,t,f_ quantify the amplitude (i.e. 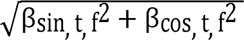) and the shape (i.e. arctan 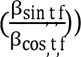) of the NBLF, that is, the causal prior’s phase modulation at a given time-frequency point. NBLFs were fitted and their amplitude and shape analysed as a function of time and frequency. Note that the remaining BCI parameters and pCommon_const_ were taken from the BCI model initially fitted across the three main experimental conditions (see above and Tab. S1) and were not optimized in the second fitting procedure. For each time-frequency point, the BADS optimization algorithm was initialised 5 times with random values for parameters β_sin,t,f_ and β_cos,t,f_ and the NBLF with the highest log likelihood was used across these initialisations. This optimization procedure resulted in reliable and accurate recovery^113^ of both β_sin,t,f_ and β_cos,t,f_ parameters (supplemental Fig. S10).

The individually fitted NBLFs enabled the prediction of the causal priors at different prestimulus time points, frequency and phases, and also enabled predictions for CMB and causal judgement (supplemental Fig. S7). The shape of the NBLFs was tested for consistency across participants using Raleigh tests for uniformity, which were complemented by Bayes factors (BF_10_) computed from a Bayesian test of circular uniformity using the ‘BayesCircIsotropy’^114^ package for R (https://github.com/keesmulder/BayesCircIsotropy). This test compared the NBLF shape to a unimodal Van Mises distribution with a non-informative Jeffreys prior (i.e. with a truncation of the concentration parameter at κ = 20).

To test whether the NBLFs predict the modulation of the causal prior by alpha phase, the NBLF amplitude (8-12 Hz) was baseline-corrected as the percent signal change relative to the average amplitude of the prestimulus baseline (i.e. -0.7 to -0.6 s before AV target onset). The NBLF amplitude was then tested against zero using a nonparametric randomisation test involving 5,000 random sign flips, based on a one-sided one-sample t-test with an auxiliary cluster-defining threshold of *t* = 2.

To further investigate the neural mechanisms underlying the NBLF amplitude (8-12 Hz, -700 to 100 ms) beyond the alpha phase, four candidate mechanisms that were accessible in the EEG data and which have been discussed in previous literature^19,74–79^, were examined: (i) Instantaneous alpha frequency, as defined above. (ii) The amplitude of occipital dipole activity (averaged between 8 and 12 Hz), computed using Fieldtrip’s^115^ dipole fitting functions. This was done by localising a single dipole to the occipital cortex using the individual’s mean ERPs, as implemented in Brainstorm using with a noise covariance matrix computed from -1.4 to -0.9 s before AV target onset. The dipole amplitude was computed as the radius of the dipole activity in spherical coordinates. (iii) Functional connectivity between early visual and early auditory regions. (iv) Functional connectivity between early visual and parietal regions. Functional connectivity was defined as phase coupling and computed using source-localised and time-frequency transformed data from the left and right V1 and V2 (as defined by the Brodmann atlas in Brainstorm), auditory regions (i.e. the left and right transverse temporal sulcus/gyrus, and the planum temporale) and the parietal cortex (the left and right superior parietal lobule, the angular gyrus and the intraparietal sulcus). Phase coupling was defined as the absolute length of the average phase angle difference between two regions across trials, as measured by inter-site phase clustering (ISPC)^107^. ISPC was averaged within the alpha band (i.e. 8-12 Hz). All four predictors were computed as relative percent signal change using a prestimulus baseline between -1.4 to -1s before AV target onset, for each frequency separately. A full regression model was computed at the group level for the analysis, regressing the NBLF amplitude against the four neural mechanisms to define the uniquely explained variance of the predictors. The uniquely explained variance was defined as the difference between the explained variance of the full regression model (i.e. R^2^_full_) and the explained variance of a reduced regression model (i.e. R^2^_red_) in which one of the four mechanisms was omitted (i.e. R^2^_unique_ = R^2^_full_ - R^2^_red_). This computation is equivalent to a semi-partial correlation^116^, but it allows one to compute the uniquely explained variance of predictors with two regressors, such as phase.

As a critical test of the causality of the entrained alpha phase on perception, the NBLFs fitted to each time-frequency point were used to predict the individual effect of phase entrainment on the causal prior in the on- and off-phase entrainment conditions. Given the entrained phases C_on,t,f_ and C_off,t,f_, where the phases are averaged across an individuals’ trials, the NBLFs predicted the causal prior for both conditions (i.e. pCommon_on_,_t,f_ and pCommon_off_,_t,f_) without any additional parameters. To assess the accuracy of these predictions, we correlated the predicted causal priors for on-phase and off-phase entrainment with the observed causal priors for both entrainment conditions across participants (i.e. two data points per participant; Tab. S2). The predicted and observed causal priors were mean-corrected for each individual before computing a Spearman rank correlation. This was done to control for individual differences in the constant causal prior, and is equivalent to a repeated-measures correlation^117^. Spearman correlations were used to obtain more robust estimates of correlations (i.e. less sensitive to outliers, e.g. from large NBLF amplitudes) than parametric correlations. Specifically, Spearman correlations were computed for NBLF-predicted causal priors averaged within the alpha band (8-12 Hz) and for frequencies below and above the alpha band (Fig. 6). We then tested whether the Spearman correlations were larger than zero in nonparametric randomisation tests (i.e. 5,000 randomisations of participants’ causal priors), using the correlations’ t values as the test statistics, with cluster-based correction and a cluster-defining threshold of *t* = 2. Finally, to investigate the relationship between causal prior modulation and CMB as well as causal judgements (Fig. 6C), we computed the Spearman correlation between the difference in the NBLF-predicted on- and off-phase causal priors and the difference in the CMB or causal judgements between entrainment conditions. This was done separately for each of the three tasks in the significant cluster (Fig. 6B; i.e. at 11 Hz and -160 ms).

### Psychophysics - Modelling of physical-behavioural link functions

Unlike the NBLF, which was fitted using both behavioural and electrophysiological data, the PLBF was fitted using only behavioural data. Specifically, we fitted the BCI model with 8 parameters of the causal prior for each entrainment phase (pCommon_1_, pCommon_2_, …, pCommon_8_) and each entrainment frequency (i.e. 5, 10, and 15 Hz). We then tested whether these parameters were modulated by the entrainment phase. The causal prior pCommon_f,D_ was fitted as a sinusoidal physical-behavioural link function (PBLF) of the entrainment phase C_i,f_ for each entrainment frequency f:

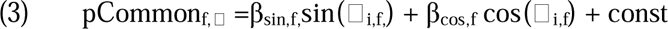

The pCommon_f,D_ parameters were mean-corrected across the 8 entrainment phases in each observer before fitting the function. The parameters β_sin,f_ and β_cos,f_ quantify the amplitude (i.e. 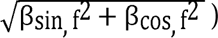) and the shape (i.e. arctan 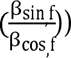) of the PBLF (i.e. the causal prior’s phase modulation) for a given entrainment frequency. Clustering of the PBLF’s shape around the circle at the group level was tested using Raleigh tests and Bayes factors. Furthermore, we tested whether the PBLF’s amplitude was larger than chance level using a nonparametric randomisation test (i.e. 5,000 randomisations of phase entrainment conditions) with a one-sided one-sample t-test as test statistic. Similarly, we compared the PBLF’s amplitudes between entrainment frequencies using a randomisation test that exchanged entrainment frequency conditions, with a two-sided paired t-test as test statistic.

## Acknowledgments

The authors thank volunteers for their participation in the experiments and Alexander Krieg and Fabian Bläsius for help with data collection in the EEG and psychophysical studies, respectively. This research was funded by a grant to TR and DS from the German Research Foundation (RO 5587/1-1, RO 5587/5-1 and SE1859/10-1).

## Data availability

All data will be made publicly available upon acceptance of the manuscript.

## Code availability

Code for computational BCI modelling will be made available to reviewers through a repository once the manuscript is sent out for review. This code will be also made publicly available upon acceptance of the manuscript.

