## Supplemental material for "Causal role of the individual alpha phase in multisensory perception"

**Supplemental methods**

*Computational modelling*

Five competing models were fitted to the behavioural data: (i) The Bayesian causal inference (BCI) model^1^ with a model-averaging strategy combines the precision-weighted fusion estimate with the auditory segregation estimate proportional to the posterior probability of a common or independent causes, respectively. (ii) The BCI model with a model-selection strategy selects the fusion or segregation estimate with the higher posterior causal probability^2^. (iii) The BCI model with a non-optimal probability-matching strategy selects the fusion/segregation estimates stochastically in proportion to the posterior causal probabilities^2^. (iv) The stochastic fusion model heuristically selects either the fusion or segregation estimate with a fixed probability that is estimated from observers’ responses^3^. (v) The fixed-criterion model heuristically selects the segregation estimate when the audiovisual spatial disparity estimate exceeds a fixed threshold^3^. Unlike the BCI model, the fixed-criterion model considers the posterior causal probabilities but applies a threshold directly on the spatial disparity estimate, ignoring observers’ causal uncertainty.

Details on the BCI model and the fitting to behavioural judgements have been described elsewhere^1,3,4^. In short, the generative model of the BCI model (Fig. 1C) assumes that common (C=1) or independent (C=2) causes are determined by sampling from a binomial distribution with a causal prior of p(C=1) = pCommon. For a common cause, the “true” number of AV target stimuli (N_AV_) is drawn from the numeric prior distribution N(μ_P_, σ_P_). For two independent causes, the “true” auditory (N_A_) and visual (N_V_) locations of the stimuli are sampled independently from the spatial prior distribution. Sensory noise is introduced by sampling the sensory inputs x_A_ and x_V_ independently from normal distributions centred on the experimentally defined number of auditory (or visual) stimuli with parameters σ_A_ (or σ_V_). The basic generative model includes different parameters: the causal prior pCommon, the spatial prior’s mean μ_P_ and standard deviation σ_P_, and the auditory and visual standard deviation σ_A_ and σ_V_, respectively. Given the sensory inputs x_A_ and x_V_, the observer infers the posterior probability of the underlying causal structure by combining the causal prior with the sensory evidence according to Bayes rule:

$$\left( 1 \right) p\left( C=1 | x\text{A},x\text{V} \right)=\frac{p(x\text{A},x\text{V}\text{|C=1})p\text{Common}}{p(x\text{A},x\text{V})}$$

The causal prior quantifies the believe of observers that the sequence of beeps and flashes arises from a common cause and should be integrated accordingly. In the case of a common cause (C=1), the optimal audiovisual numeric estimate ($\hat{S}\text{AV,C=1}$) is obtained by combining the auditory and visual numeric inputs as well as the numeric prior, which is weighted by their relative precision (i.e. a precision-weighted fusion estimate):

$$\left( 2 \right) \hat{N}\text{AV,C=1}=\frac{\frac{x\text{A}}{\sigma\text{A}\text{2}}+\frac{x\text{V}}{\sigma\text{V}\text{2}}+\frac{\mu\text{P}}{\sigma\text{P}\text{2}}}{\frac{1}{\sigma\text{A}\text{2}}+\frac{1}{\sigma\text{V}\text{2}}+\frac{1}{\sigma\text{P}\text{2}}}$$

In the case of independent causes (C=2), the optimal numeric estimates of the unisensory auditory ($\hat{N}\text{A,C=2}$) and visual ($\hat{N}\text{V,C=2}$) stimuli are independent (i.e. segregation estimates):

$$\left( 3 \right) \hat{N}\text{A,C=2}=\frac{\frac{x\text{A}}{\sigma\text{A}\text{2}}+\frac{\mu\text{P}}{\sigma\text{P}\text{2}}}{\frac{1}{\sigma\text{A}\text{2}}+\frac{1}{\sigma\text{P}\text{2}}}, \hat{N}\text{V,C=2}=\frac{\frac{x\text{V}}{\sigma\text{V}\text{2}}+\frac{\mu\text{P}}{\sigma\text{P}\text{2}}}{\frac{1}{\sigma\text{V}\text{2}}+\frac{1}{\sigma\text{P}\text{2}}}$$

Crucially, the observers do not know whether the stimuli come from common or independent sources, but they must infer this from the sensory inputs. To account for the observers’ causal uncertainty, the model computes a final task-relevant spatial estimate (e.g. $\hat{N}\text{A}$ for auditory report) by combining the fusion estimate (i.e. $\hat{N}\text{AV,C=1}$ for C=1) and the task-relevant segregation estimate (e.g. $\hat{N}\text{A,C=2}$ for C=2) depending on the posterior probabilities of the estimates’ underlying causal structures ($p\left( C=1 | x\text{A},x\text{V} \right)$. Thus, the observers report different task-relevant estimates for an auditory ($\hat{N}\text{A}$) or visual ($\hat{N}\text{V}$) report. In the ‘model averaging’ strategy of the BCI model, the observers weigh the estimates in proportion to the posterior probabilities of their underlying causal structures:

$$\left( 4 \right) \hat{N}\text{A }\text{= p(C=1|}x\text{A},x\text{V}\text{)} \hat{N}\text{AV,C=1 }\text{+ (1 - p(C=1|}x\text{A},x\text{V}\text{)} )\hat{N}\text{A,C=2 }\text{ }$$

$$\hat{N}\text{V }\text{= p(C=1|}x\text{A},x\text{V}\text{)} \hat{N}\text{AV,C=1 }\text{+ (1 - p(C=1|}x\text{A},x\text{V}\text{)} )\hat{N}\text{V,C=2}$$

In the model selection strategy, the observers select the numeric estimate that has a higher posterior probability:

$$\left( 5 \right) \hat{N}\text{A }\text{= }\left\{ \begin{aligned} \hat{N}\text{A,C=1 }\text{if}\text{ }\text{p(C=1|}x\text{A},x\text{V}\text{)}> 0.5 \\ \hat{N}\text{A,C=2 }\text{if}\text{ }\text{p(C=1|}x\text{A},x\text{V}\text{)}\leq0.5 \end{aligned} \right. , \hat{N}\text{V }\text{= }\left\{ \begin{aligned} \hat{N}\text{V,C=1 }\text{if}\text{ }\text{p(C=1|}x\text{A},x\text{V}\text{)}> 0.5 \\ \hat{N}\text{V,C=2 }\text{if}\text{ }\text{p(C=1|}x\text{A},x\text{V}\text{)}\leq0.5 \end{aligned} \right.$$

In probability matching, the observers select the numeric estimate stochastically in proportion to the posterior causal probabilities:

$$\left( 6 \right) \hat{N}\text{A }\text{= }\left\{ \begin{aligned} \hat{N}\text{AV,C=1}\text{ if}\text{ }\text{p(C=1|}x\text{A},x\text{V}\text{)}> \alpha\\ \hat{N}\text{A,C=2}\text{ if}\text{ }\text{p(C=1|}x\text{A},x\text{V}\text{)}\leq\alpha\end{aligned} \right., \hat{N}\text{V }\text{= }\left\{ \begin{aligned} \hat{N}\text{AV,C=1}\text{ if}\text{ }\text{p(C=1|}x\text{A},x\text{V}\text{)}> \alpha\\ \hat{N}\text{V,C=2}\text{ if}\text{ }\text{p(C=1|}x\text{A},x\text{V}\text{)}\leq\alpha\end{aligned}, \alpha\sim U\left( 0,1 \right) \right.$$

These three Bayesian decision strategies that take causal uncertainty into account were compared with two additional non-optimal heuristic strategies^3^: In the stochastic-fusion model, observers stochastically choose the audiovisual numeric (i.e. fusion) or the task-relevant unisensory auditory or visual (i.e. segregation) estimate with a fixed probability parameter η:

$$\left( 7 \right) \hat{N}\text{A }\text{= }\left\{ \begin{aligned} \hat{N}\text{AV,C=1}\text{ if }\eta>\alpha\\ \hat{N}\text{A,C=2}\text{ if}\text{ }\eta\leq\alpha\end{aligned} \right., \hat{N}\text{V }\text{= }\left\{ \begin{aligned} \hat{N}\text{AV,C=1}\text{ if}\text{ }\eta>\alpha\\ \hat{N}\text{V,C=2}\text{ if}\text{ }\eta\leq\alpha\end{aligned}, \alpha\sim U\left( 0,1 \right) \right.$$

The stochastic-fusion model encompasses a potential ‘forced’ fusion (η = 1) or ‘forced’ segregation (η = 0) as well as any intermediate response behaviour (0 < η < 1). The fixed-criterion model reports a fusion or segregation estimate when the absolute spatial disparity of the stimuli is below or above a fixed criterion parameter k. Unlike the BCI model, the fixed-criterion model implements a response heuristic based on audiovisual disparity without formally computing the posterior probability of the causal structure:

$$\left( 8 \right) \hat{N}\text{A }\text{= }\left\{ \begin{aligned} \hat{N}\text{AV,C=1}\text{ if }\left| x\text{A}-x\text{V} \right|<k \\ \hat{N}\text{A,C=2}\text{ if }\left| x\text{A}-x\text{V} \right|\geq k \end{aligned} \right., \hat{N}\text{V }\text{= }\left\{ \begin{aligned} \hat{N}\text{AV,C=1}\text{ if }\left| x\text{A}-x\text{V} \right|<k \\ \hat{N}\text{V,C=2}\text{ if }\left| x\text{A}-x\text{V} \right|\geq k \end{aligned} \right.$$

For the causal judgement task, it was assumed that participants reported a common source when the posterior probability of a common source was greater than the threshold of 0.5 ^5^.

$$\left( 9 \right) \hat{C}\text{ }\text{= }\left\{ \begin{aligned} \text{1 if}\text{ }\text{p(C=1|}x\text{A},x\text{V}\text{)}>0.5 \\ \text{2 if}\text{ }\text{p(C=1|}x\text{A},x\text{V}\text{)}\leq0.5 \end{aligned} \right.$$

To compare between the five candidate models, each model was fitted to the participants’ visual and auditory numerical reports, as well as their causal judgements across in AV conditions in both studies. For model fitting, the predicted distributions of the auditory and visual numeric estimates (i.e. the marginal distributions: p($\hat{N}\text{A}\text{|N}\text{A}\text{, N}\text{V}$) and p($\hat{N}\text{V}\text{|N}\text{A}\text{, N}\text{V}$) and the posterior probability of a common cause (i.e. p($\hat{C}\text{|N}\text{A}\text{, N}\text{V}$)) were used. These distributions were obtained by marginalizing over the internal variables x_A_ and x_V_ that were not accessible to the experimenter^1^. These distributions were generated by simulating x_A_ and x_V_ for each experimental condition (i.e. continuous variables sampled from Gaussian distributions) 5,000 times for the EEG study or 2,500 times for the psychophysical study, and inferring $\hat{N}\text{A}$ and $\hat{N}\text{V}$ as well as $\hat{C}$ from equations (1) through (9). For the auditory and visual numerical reports, the predicted distributions of $\hat{N}\text{A}$ and $\hat{N}\text{V}$ were linked to participants’ categorical numerical reports (i.e. four possible signal numbers) by assuming that participants selected the button that is closest to $\hat{N}\text{A}$ / $\hat{N}\text{V}$. $\hat{N}\text{A}$ / $\hat{N}\text{V}$ were binned accordingly into a four-bin histogram. From these predicted multinomial distributions (i.e. one for each of the experimental conditions), the log likelihood of the numerical reports was summed, given the predicted distributions across all AV conditions with numerical reports (i.e. 96 AV conditions in the EEG study and 768 AV conditions in the psychophysical study). For the causal judgement task, the log likelihoods of the binary causal judgements were summed, given the predicted distributions for $\hat{C}$ across the AV conditions of this task (i.e. 32 AV conditions in the EEG study and 384 AV conditions in the psychophysical). Finally, the log likelihoods were summed across the numerical reports and causal judgements. Please note that in the EEG study the competing models were fitted across the three experimental entrainment conditions (i.e., control, on-phase and off-phase) as well as within each of the conditions (Tab. S2).

A Bayesian optimization algorithm as implemented in the BADS toolbox^6^ was used to obtain maximum likelihood estimates for the models’ parameters. To identify the optimal model for explaining the participants’ data, the five candidate models were compared using the Bayesian information criterion (BIC) as an approximation to the model evidence^7^. Bayesian model comparison^8^ was performed at the random-effects group level as implemented in SPM12^9^ to obtain the protected exceedance probability (i.e. the probability that a given model is more likely than any other model, beyond differences due to chance^8^) for each of the four candidate models. Next, predictions for the behavioural CMB and causal judgements were generated based on the fitted BCI model’s prediction in the EEG study (Fig. 2A, B). Thus, new x_A_ and x_V_ values were simulated for 10,000 trials for each experimental condition using the fitted model parameters of each participant. Finally, the kernel-density estimated maximum-a-posteriori estimates of p($\hat{N}\text{A}\text{|N}\text{A}\text{, N}\text{V}$) and p($\hat{N}\text{V}\text{|N}\text{A}\text{, N}\text{V}$) and p($\hat{C}\text{|N}\text{A}\text{, N}\text{V}$) were analysed for each condition, exactly as done for participants’ numerical reports and causal judgements.

**Supplemental results**

*Multisensory integration and entrainment effects on behavioural indices*

The crossmodal bias (CMB), which quantifies the relative influence of auditory and visual signals on numerical reports^5,10^, replicated the key profile of multisensory integration when observers performed causal inferences^4,5,11^. The number of beeps strongly biased the visual reports, particularly when the numeric disparity between flashes and beeps was small, indicating a common cause (Fig. 2A; see supplemental Fig. S1 for individual results). In contrast, the number of flashes had only a weak influence on the auditory reports (i.e. interaction of task relevance x numeric disparity, F_1.6, 38.8_ = 132.244, p < 0.001, part. η^2^ = 0.841, BF_10_ > 100; supplemental Tab. S1). Overall, the control condition had stronger visual influences on the CMB than the entrainment conditions (main effect of experimental entrainment condition, F_1.4,35.4_ = 15.525, p < 0.001, part. η^2^ = 0.383, BF_10_ > 100; see supplemental results and Fig. S2). This result could be related to higher visual uncertainty from the flickering visual entraining stimuli in the on-phase and off-phase conditions (see below). Most importantly, the key profile of CMB was equivalent for both on-phase and off-phase entrainment (main effect of on-phase vs. off-phase, F_1,25_ = 0.250, p = 0.621, part. η^2^ = 0.010, BF_10_ = 0.149). For causal judgements, the proportion of common-cause judgements decreased as the numeric disparity increased (main effect of numeric disparity, F_1.5,37.9_ = 165.466, p < 0.001, part. η^2^ = 0.869, BF_10_ > 100; Tab. S1). In the control condition, participants more often reported a common cause for congruent AV signals (i.e. interaction effect of experimental entrainment condition and numeric disparity; F_4.2,104.9_ = 5.274, p < 0.001, part. η^2^ = 0.174, BF_10_ > 100). Critically, Bayes factors revealed substantial evidence that the causal judgements were equivalent in the on-phase and off-phase entrainment conditions (main effect of on-phase vs. on-phase, F_1, 25_ = 0.001, p = 0.970, part. η^2^ < 0.001, BF_10_ = 0.141).

*Model comparison and analysis of BCI model parameters*

Using Bayesian model comparisons (see Tab. S2), we found that the BCI model with model averaging as the causal decision strategy clearly outperformed the four competing computational models with alternative Bayesian causal decision strategies (i.e. model selection or probability matching) or heuristic decision strategies (i.e. stochastic fusion or fixed criterion). This pattern was found when fitting the models across the three experimental entrainment conditions as well as within each of the conditions. Thus, participants consistently applied the same causal decision strategy in all three entrainment conditions in the EEG study. This finding from the EEG study was replicated in the psychophysical study (Tab. S4), and it aligns with our previous results for the BCI model in the sound-induced flash illusion paradigm^4,12^.

When we compared the BCI model parameters between the two entrainment conditions (Fig. 2C, Fig. S3 and Tab. S3), we found evidence of equivalence for the causal prior (BF_10_ = 0.209), the numeric prior (BF_10_ = 0.208) and the auditory variance (BF_10_ = 0.253) parameters. The remaining two parameters, i.e. the numeric prior’s mean and the visual variance, did not differ significantly between the two conditions (p > 0.05), even though Bayes factors indicated inconclusive evidence (i.e. BF_10_ = 0.778 for μ_P_ and BF_10_ = 1.215 for σ_V_).

When comparing the two entrainment conditions to the control condition, we observed that the mean of the numeric prior and the visual variance were significantly higher in the entrainment conditions. However, these differences could be attributed to the flickering visual stimuli in the entrainment conditions and were unrelated to variations in the causal prior. Interestingly, participants occasionally counted the final stimulus of the entraining flickering visual stimuli as an additional target flash. This is evident in the higher number of flashes and greater visual uncertainty in the entrainment conditions compared to the control condition (supplemental Fig. S2) in both in AV and unisensory conditions with visual report. The larger visual variance (i.e. higher visual uncertainty) in the entrainment conditions could also explain the stronger auditory influence on the CMB in these conditions (Fig. 2A), as visual signals receive smaller weights during integration. This is consistent with accounts of precision-weighted integration^13,14^. Specifically, the BCI model averages the audiovisual signals weighted by their relative precision (i.e. the inverse of variance; equation (2)), in the case of a common cause. Thus, larger visual variance parameters lead to a stronger relative auditory influence on the model’s CMB predictions, as was indeed found for the CMB in the onphase and off-phase conditions (Fig. 2A).

*Behavioural consequences of NBLFs with opposite shapes*

When two individuals have NBLFs with nearly opposite shapes, the NBLFs predict opposite entrainment effects on the causal prior (Fig. 5B). These opposite effects translate into opposite effects on multisensory interactions, as demonstrated by the NBLFs prediction for CMB and causal judgements. Supplemental figure S6 shows that opposite NBLF shapes lead to reversed effects of on-phase versus off-phase entrainment on the CMB and causal judgements for the two observers depicted in Fig. 5B. This explains why, when averaged at the group level, the CMB and causal judgements are equivalent for the two entrainment conditions (see Fig. 2A), especially when the NBLFs are inconsistent between observers.

*Rationale for modelling individual NBLFs in entrainment studies*

If the relationships between alpha phase and behaviour are nonlinear (e.g. sinusoidal) and unique to each individual, then the specific mapping between phase and behaviour must be known in order to test whether the entrained alpha phase causes a perceptual effect. Only with such explicit mappings can one exactly predict how a manipulated phase will affect the causal prior according to the preferred or non-preferred phases (i.e. the NBLF’s shape; supplemental Fig. S9). Even in the case of a generic NBLF, the entraining stimuli can lead to a positive or negative difference, or even no difference (i.e. in the unlikely case that entrained phase exactly hits the function’s two inflection points) in the causal prior between the entrainment conditions, depending on the NBLF’s specific shape. Individual NBLFs exacerbate this fundamental challenge because they result in a wide range of positive and negative entrainment effects on the causal prior, with an expected value of zero at group level (supplemental Fig. S9 and Fig. 1E). In other words, generic NBLFs may lead to positive or negative entrainment effects of two opposite phase manipulations. However, individual phase-behaviour relationships, which have been often been demonstrated in perception^4,15–21^, will inevitably lead to a null effect at group level. Essentially, the simulation in figure S9 shows that the severity of ignoring individual NBLFs depends on the consistency of NBLFs shapes in the sample. For example, this can be modelled by the concentration parameter of a circular van Mises distribution from which shape is sampled. The lower the consistency of NBLF shape, the higher is the probability of missing a phase-behaviour relationship at the group level. Therefore, modelling NBLFs is essential for predicting the individual quantitative size of the alpha-phase entrainment effect on the causal prior (Fig. 1E). Since this approach has not yet been implemented, its absence could explain the heterogeneity of findings in causal alpha-phase entrainment studies.

*Psychophysical study*

In the psychophysical study, we analysed the generic effects of the eight phase steps on the causal prior using repeated-measures ANOVAs. The main effects of phase were not significant and the causal priors were equivalent across the phase steps (5 Hz: F_4.8,136.9_ = 0.855, p = 0.511, part. η^2^ = 0.030, BF_Incl_ = 0.039; 10 Hz: F_7,196_ = 0.774, p = 0.610, part. η^2^ = 0.027, BF_Incl_ = 0.032; 15 Hz: F_7,196_ = 1.00, p = 0.433, part. η^2^ = 0.034, BF_Incl_ = 0.049).

**Supplemental figures**

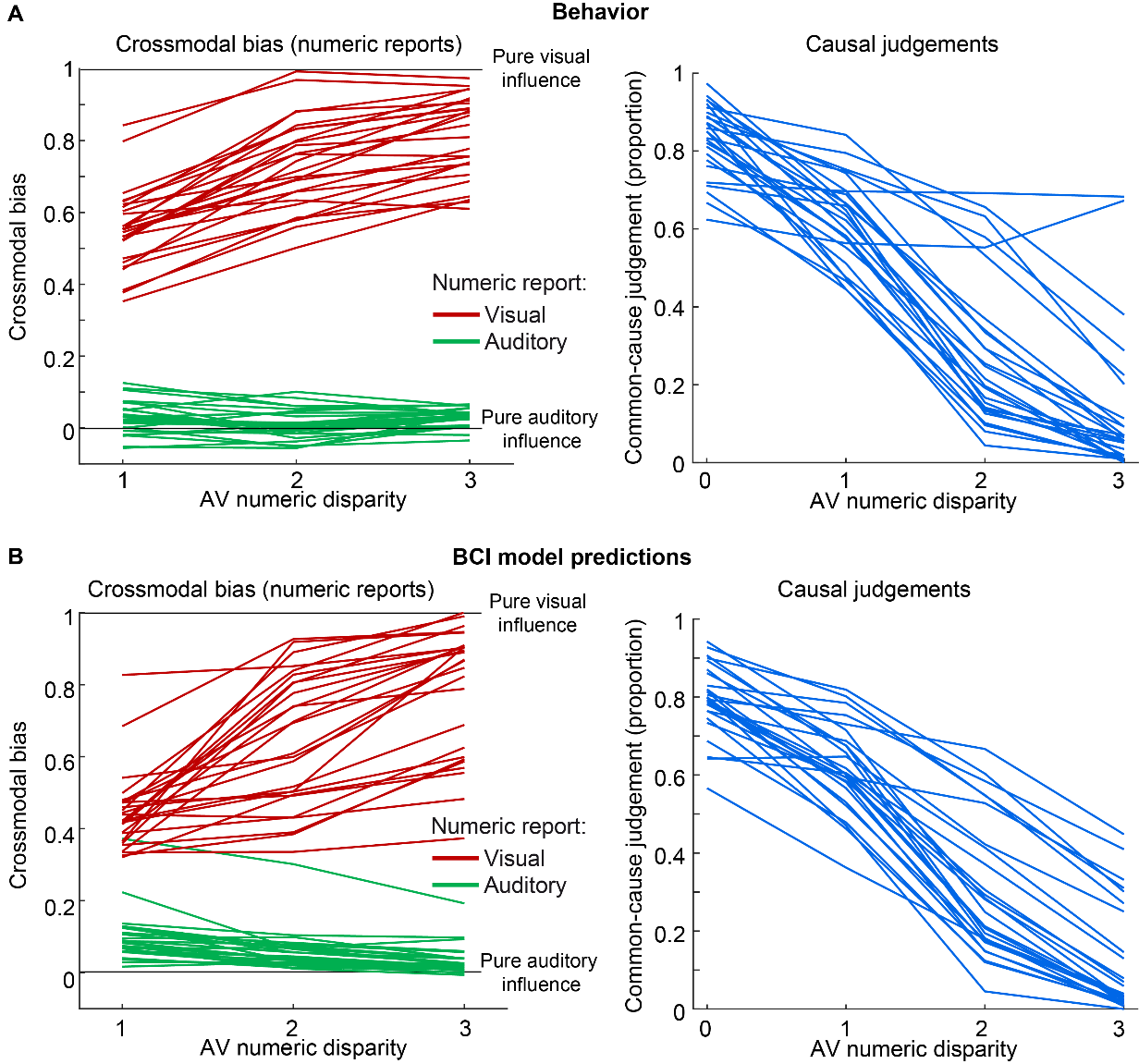

**Figure S1. Individual behavioural data show consistent patterns of crossmodal bias (CMB) and causal judgements as predicted by the BCI model. (A)** CMB and causal judgements computed from participants’ reports. **(B)** CMB and causal judgements computed from the BCI model’s predictions (i.e., model averaging).

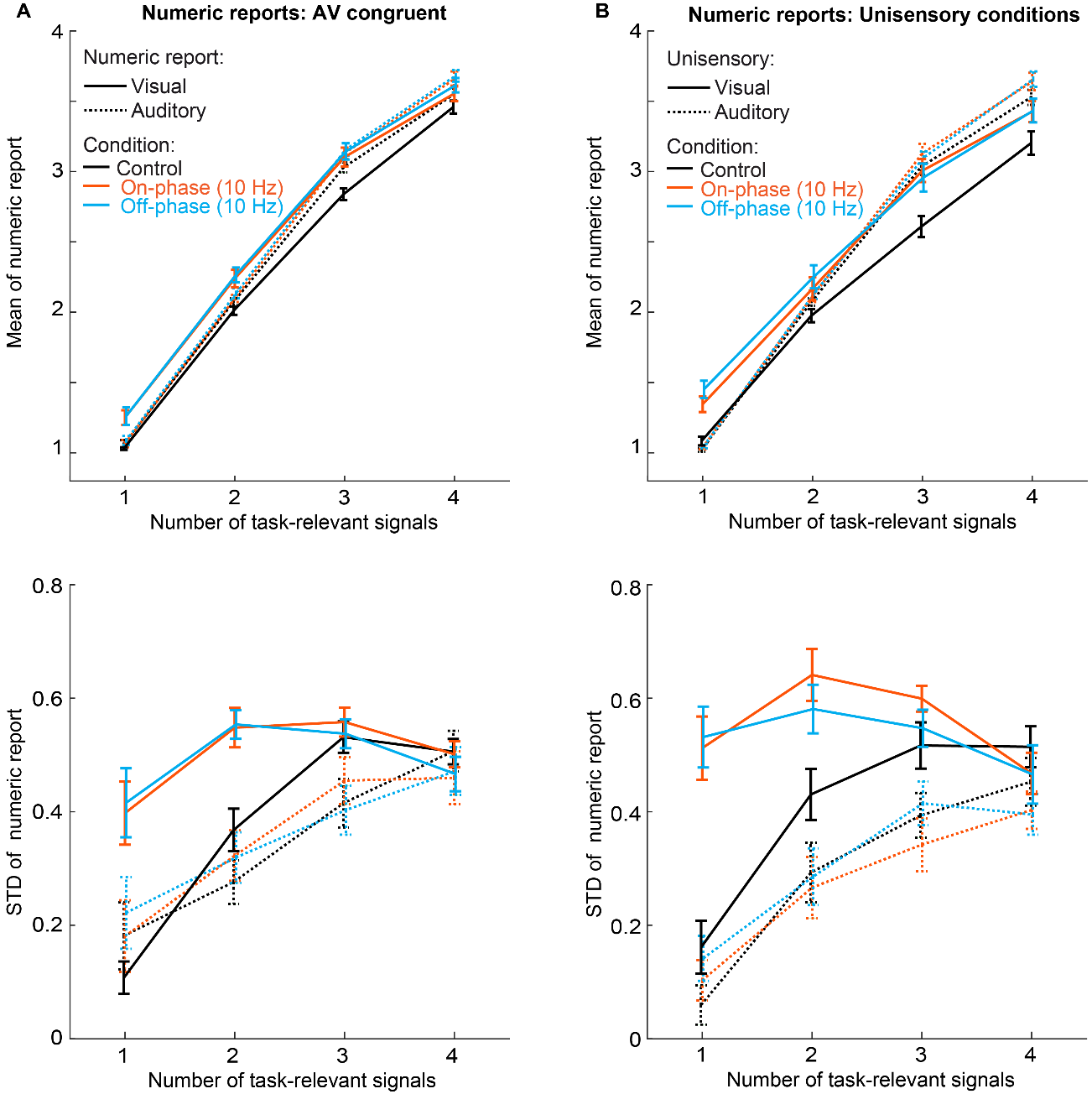

**Figure S2. The on-phase and off-phase entrainment conditions increase the number of perceived flashes and the uncertainty of visual numerical reports compared to the control condition.** The larger visual variance in the on-phase and off-phase conditions explains the stronger auditory influence on the CMB in these conditions (Fig. 2A), as visual signals receive smaller weights during integration. Plots show the mean and STD of numerical reports (across-participants mean ± SEM) in AV congruent (i.e. same number of visual and auditory stimuli) and unisensory conditions as a function of task-relevant signal number, for V and A report as well as the three entrainment conditions. **(A)** Mean and STD of numerical reports in AV congruent conditions. **(B)** Mean and STD of numerical reports in unisensory auditory and unisensory visual conditions.

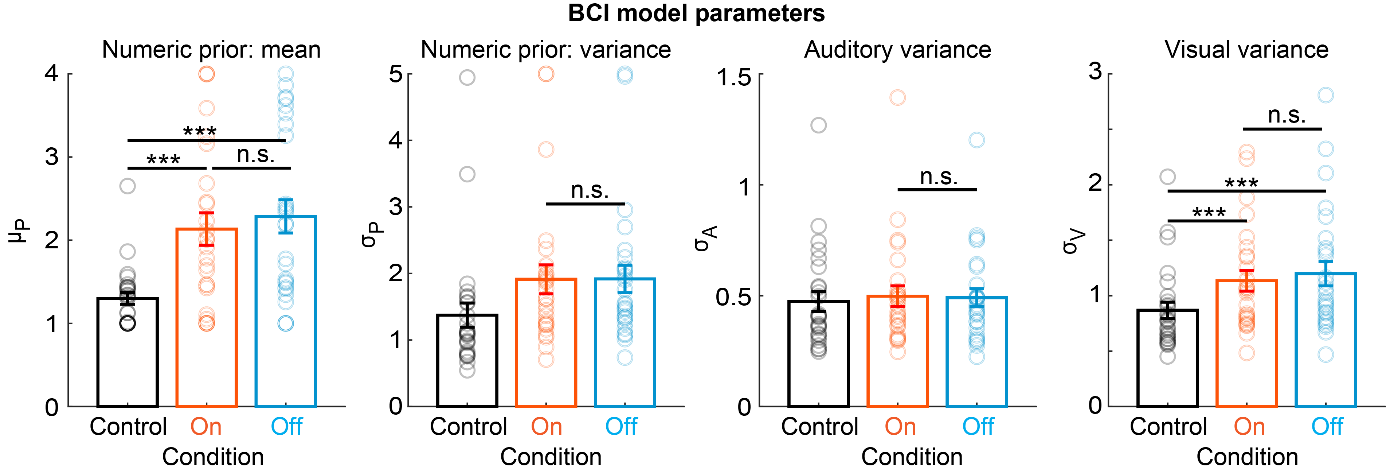

**Figure S3. The BCI models numeric prior and visual/auditory variance parameters show no differences between on-phase and off-phase entrainment conditions.** The parameters of the BCI model are plotted separately for each of the three experimental conditions (for the causal prior parameter, see Fig. 2C). The BCI model’s decision strategy applies model averaging. Significant differences are indicated by * = p < 0.05 and *** < 0.001. Non-significant differences are indicated by “n.s.”.

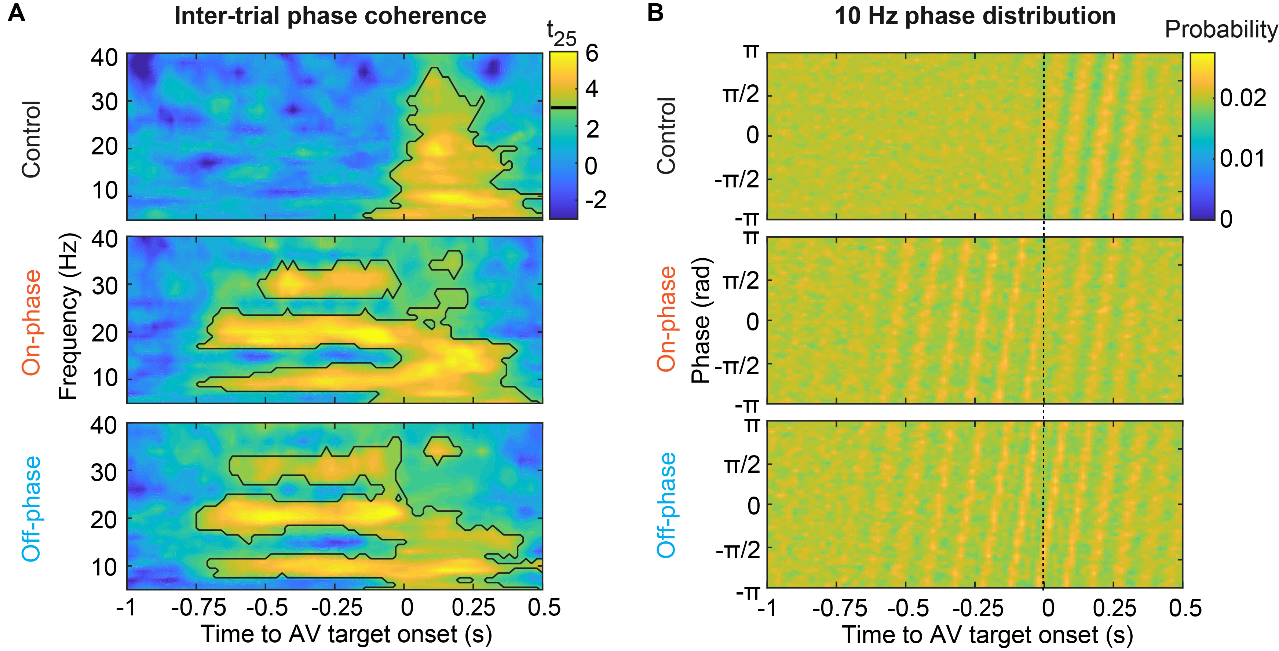

**Figure S4. On-phase and off-phase 10 Hz visual stimulation evoke phase-locked 10 Hz oscillations with opposite phase angles in early visual areas.** Inter-trial phase clustering (ITPC) and phase distribution were computed from source-localised data in V1 and V2. **(A)** Time-frequency t-value maps of ITPC for the three entrainment conditions. Significant clusters (p < 0.001; two-sided cluster-based-corrected randomisation t_25_ test; cluster-defining threshold t = 3) are demarcated by a solid line. **(B)** Distributions of phase (across-participants mean) at 10 Hz.

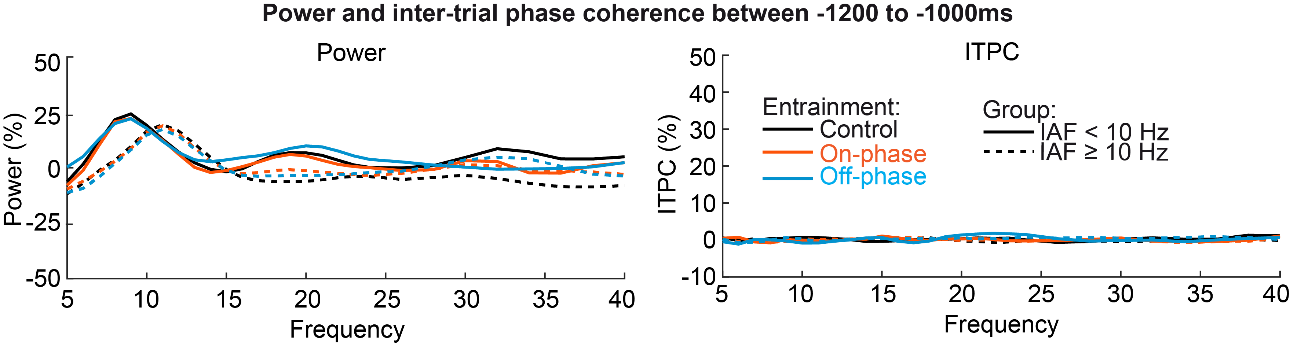
 **Figure S5. Before the onset of entraining stimuli, power and inter-trial phase coherence show power peaks at the individual alpha frequency, but entrainment has not yet occurred.** Power and intertrial-phase coherence (ITPC) (across-participants mean; normalized relative to baseline) plotted as a function of frequency and individual alpha frequency (IAF; measured from resting-state data) below (n = 10; 9.294 +- 0.359 Hz; mean +- STD IAF) and above 10 Hz (n = 16; 10.821 ± 0.786 Hz IAF). ITPC is averaged for a time window from -1200 to -1000ms before onset of the entraining stimuli, which occurred -933 and -533 ms before AV target onset.

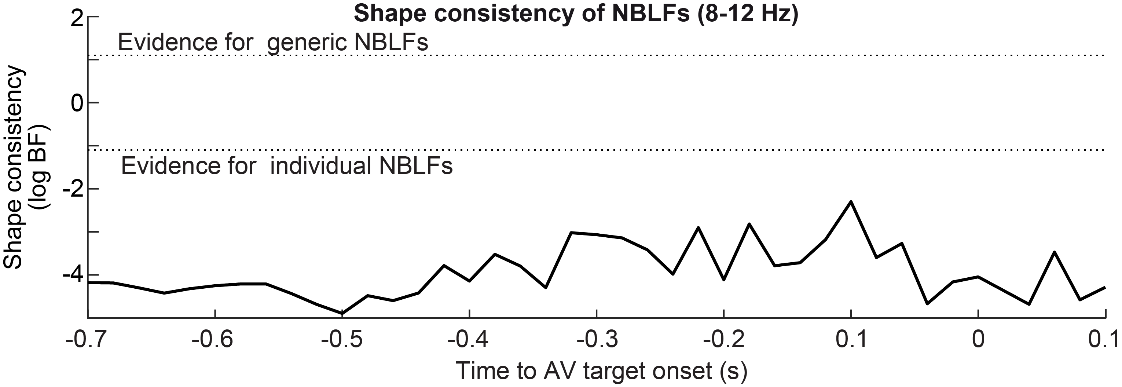

**Figure S6. NBLF shape in the alpha band (8-12 Hz) is inconsistent across individuals.** Bayes factors (logBF) quantify the relative evidence for inconsistent individual (i.e. uniformly distributed) versus consistent generic (i.e. unimodally distributed) shapes of NBLFs for alpha band oscillations (8-12 Hz). Bayes factors provided strong to decisive empirical support for individual NBLFs throughout the prestimulus time window. Bayes factors were computed from a Bayesian test of circular uniformity.

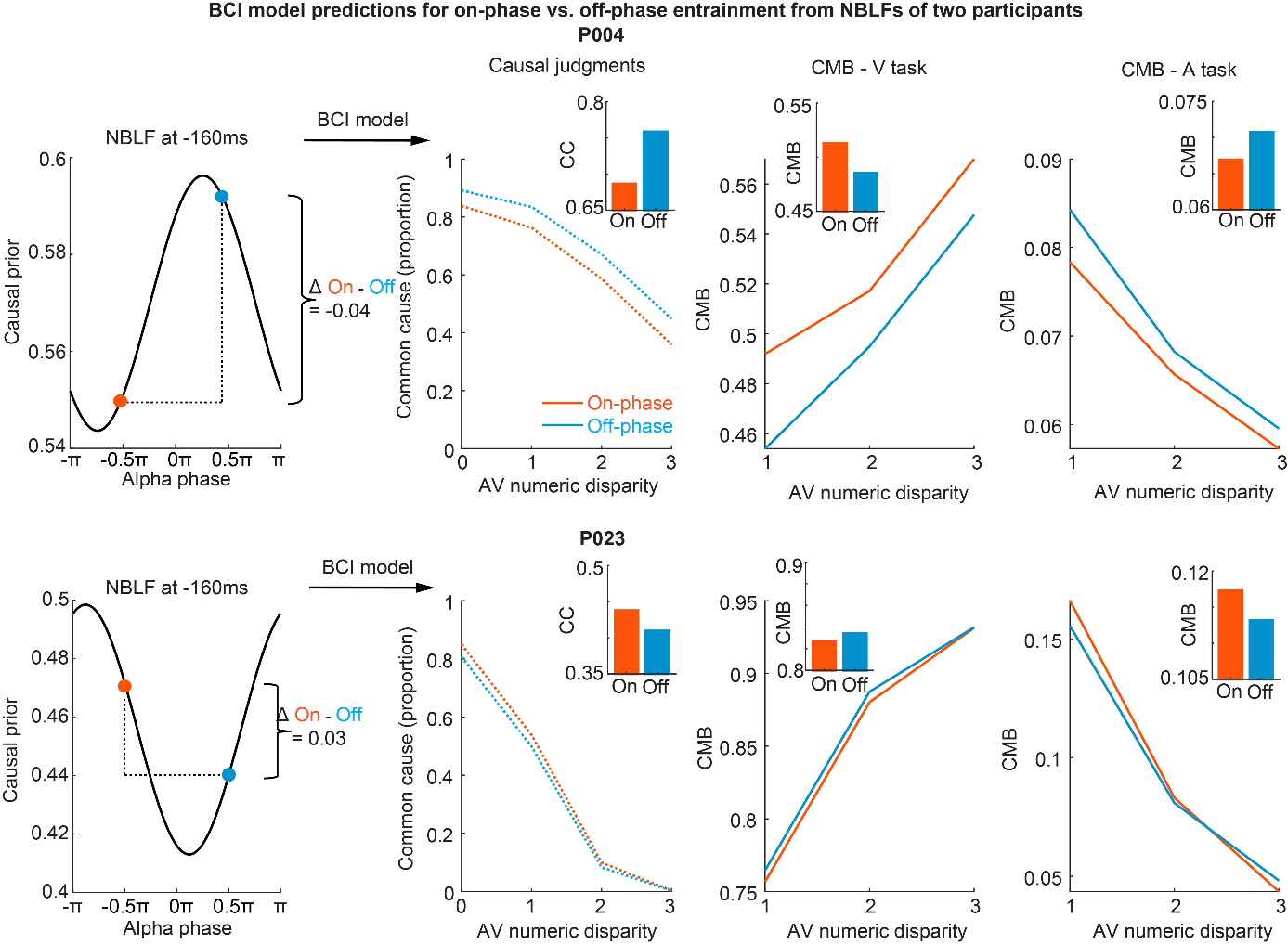

**Figure S7. The BCI model predicts opposite effects for CMB and causal judgements for two participants (P004 and P023) with two opposite NBLFs.** The full NBLFs are shown in Fig. 5B. Left panels: NBLFs were fitted to the prestimulus alpha phases over visual cortex in the control condition, here shown for -160 ms before the onset of AV target stimuli. The NBLFs predict opposite effects of the entrained alpha phase on the causal prior in both individuals for similar alpha phases. Middle and right panels: Because the NBLFs model opposite modulations of the causal prior, the BCI model predicts opposite effects of alpha phase entrainment on common-cause judgements (CC) and crossmodal bias (CMB) for the two individuals. Insets show the main effect of on-phase vs. off-phase entrainment, averaged over AV numeric disparity.

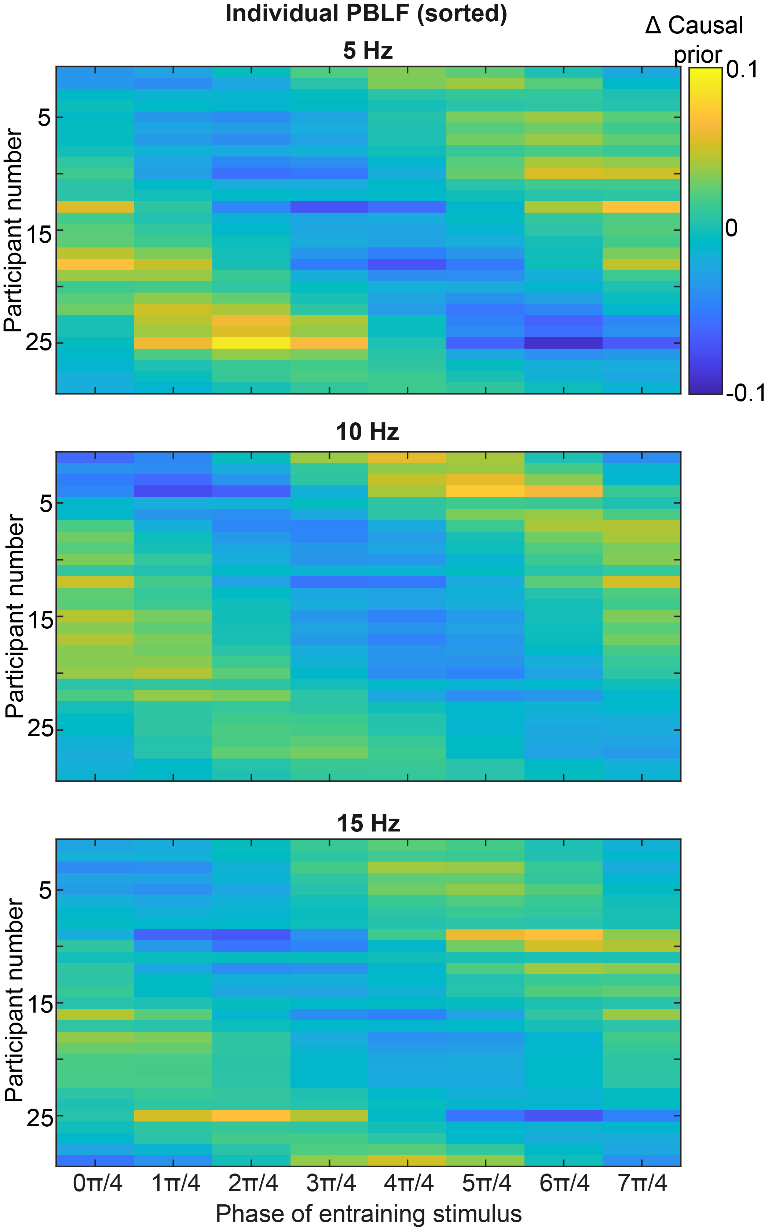

**Figure S8. Individual PBLFs in the psychophysical study.** Individual PBLFs were sorted vertically by their shape across the sample, in relation to the phase of the entraining visual stimulus (x-axis). The shapes of the PBLFs varied among individuals for all three entraining frequencies (Fig. 7D). However, the amplitudes of the PBLFs were stronger at 5 and 10 Hz (Fig. 7E), as indicated by the more saturated colours, than at 15 Hz.

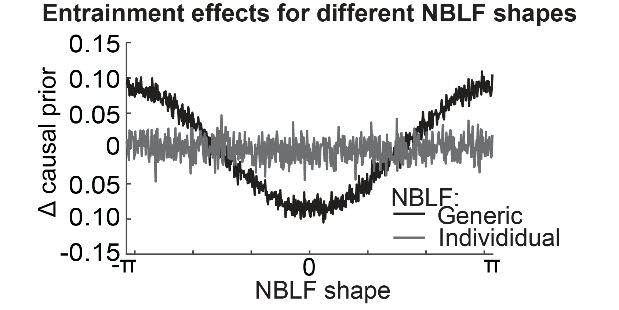

**Figure S9. Simulations show that alpha-phase entrainment effects on the causal prior depend on the shape and generality of NBLFs.** The direction and size of entrainment effects on the group-level Δ causal prior (i.e. group-average of individual difference of causal prior between on-phase versus off-phase entrainment; assuming two opposite (180°) phases of entrainment). Entrainment effects are shown for generic and individual NBLFs: For generic NBLFs, entrainment effects at group level exist and depend on the specific shape of the generic NBLFs. For individual NBLFs, no entrainment effect at group level exists because entrainment effects predicted by individual NBLFs cancel out at group level. Thus, if the NBLFs that model individual alpha phase-perception relationships are not explicitly considered, existing alpha-phase effects at the individual level will be missed when averaging data across observers, leading to false-negative group-level results. In this figure, entrainment effects were computed from the NBLFs of 26 simulated participants, where the NBLF parameters (i.e. amplitude and shape) were sampled. The NBLFs’ amplitude was sampled from a Gaussian distribution (μ = 0.05, σ = 0.02) in the same way for generic and individual NBLFs. The mean of a van Mises distribution, from which the NBLFs’ shape was sampled, systematically varied between -π to + π. (i.e. x axis). The distinction between generic and individual NBLFs was implemented by the concentration parameter of the van Mises distribution: For generic NBLFs, the simulation assumed a near-fixed effect of shape (i.e. shape distribution concentrated at a specific phase angle according to a van Mises distribution with κ = 0.1). For individual NBLFs, the simulation assumed a near-random effect of shape (i.e. nearly uniform shape distributed with κ = 4).

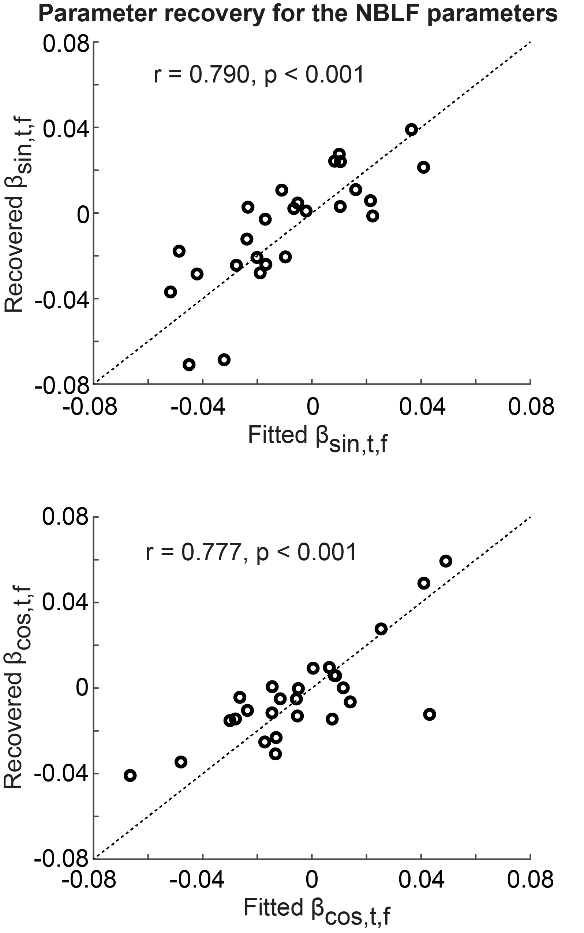

**Figure S10. Simulations demonstrate accurate and reliable parameter recovery for the NBLF’s parameters (β_sin,t,f_ and β_cos,t,f_).** The plots show the recovered parameters as a function of the parameters originally fitted to participants’ behavioural data. Note that the NBLF’s amplitude and shape parameters can be directly computed from those (see Methods). For parameter recovery, the winning BCI model (i.e. model averaging) first predicted responses based on participants’ fitted parameters at 11 Hz at -200 ms (n.b.: we used 6 replications of the data to reduce simulation noise). Next, the predicted responses were fitted to obtain recovered parameters with the same fitting procedure as for the main analysis (i.e. initialization with 5 different random parameters; predicted distributions were generated from 500 simulated trials per condition). The dotted line is a line with slope 1 and intercept 0.

**Supplemental tables**

**Table S1**

| **Table S1. Main and interaction effects of task relevance (TR), alpha entrainment (AE) and AV numeric disparity (ND) on the crossmodal bias and the causal judgements (n = 26) from repeated-measures ANOVAs.** | | | | | |
| --- | --- | --- | --- | --- | --- |
| **Crossmodal bias** | **F** | **df1, df2** | **p** | **part. η^2^** | **BF_incl_** |
| TR | 780.025 | 1, 25 | <0.001 | 0.969 | >100 |
| AE | 15.525 | 1.4, 35.4 | <0.001 | 0.383 | >100 |
| AE* | 0.250 | 1, 25 | 0.621 | 0.010 | 0.149 |
| ND | 102.453 | 1.5, 37.7 | <0.001 | 0.804 | >100 |
| TR x AE | 1.091 | 1.4, 33.8 | 0.324 | 0.042 | 0.551 |
| TR x ND | 132.244 | 1.6, 38.8 | <0.001 | 0.841 | >100 |
| AE x ND | 1.347 | 3.0, 73.8 | 0.266 | 0.051 | 0.225 |
| TR x AE x ND | 0.859 | 2.7, 67.1 | 0.456 | 0.033 | 0.036 |
| Causal judgements |  |  |  |  |  |
| AE | 3.475 | 1.4, 35.9 | 0.056 | 0.122 | >100 |
| AE* | 0.001 | 1, 25 | 0.970 | <0.001 | 0.141 |
| ND | 165.466 | 1.5, 37.9 | <0.001 | 0.869 | >100 |
| AE x ND | 5.274 | 4.2, 104.9 | <0.001 | 0.174 | >100 |
| AE* x ND | 0.197 | 2.3, 57.3 | 0.850 | 0.008 | 0.044 |
| Note: The factor AE comprised all three levels of alpha entrainment (control, on-phase, off-phase). BF_incl_ were computed from Bayesian repeated-measures ANOVAs with multivariate Cauchy priors on the effects and uniform model priors. AE* and AE* x ND is computed from a repeated-measures ANOVA which included only two levels of the factor alpha entrainments, on- vs. off-phase. For these repeated-meausres ANOVAs, all remaining main and interactions effects were highly similar to those reported above. Effects were Greenhouse-Geisser corrected if sphericity was violated. | | | | | |

**Table S2**

| **Table S2. Results of the Bayesian model comparison in the EEG study. The BCI model with three decision strategies (model averaging, model selection and probability matching) and heuristic models (stochastic fusion and fixed threshold) were compared. The models were fitted either separately for each of the three alpha entrainment conditions, or across all condition.** | | | | | | | | | | |
| --- | --- | --- | --- | --- | --- | --- | --- | --- | --- | --- |
| **Model name** | **Alpha entrainment** | **_p_Common**  η  k | **µ_P_** | **σ_P_** | **σ_A_** | **σ_V_** | **R^2^** | **relBIC** | **pEP** | **% win** |
| Model averaging | All | 0.48±0.02 | 1.70±0.14 | 1.82±0.21 | 0.50±0.04 | 1.04±0.08 | 0.85±0.01 | 6298 | 1 | 0.65 |
|  | Control | 0.52±0.01 | 1.30±0.07 | 1.37±0.18 | 0.47±0.04 | 0.87±0.07 | 0.84±0.01 | 3254 | 1 | 0.65 |
|  | On-phase | 0.48±0.02 | 2.13±0.20 | 1.91±0.22 | 0.50±0.05 | 1.13±0.09 | 0.79±0.02 | 1625 | 0.94 | 0.54 |
|  | Off-phase | 0.48±0.02 | 2.29±0.20 | 1.92±0.21 | 0.49±0.04 | 1.20±0.11 | 0.79±0.02 | 1574 | 0.95 | 0.5 |
| Model selection | All | 0.50±0.02 | 1.81±0.13 | 1.51±0.21 | 0.52±0.04 | 1.01±0.09 | 0.84±0.01 | 5801 | 0 | 0.12 |
|  | Control | 0.53±0.01 | 1.50±0.07 | 1.12±0.12 | 0.48±0.04 | 0.88±0.08 | 0.84±0.01 | 3034 | 0 | 0.19 |
|  | On-phase | 0.49±0.02 | 2.05±0.15 | 1.53±0.18 | 0.52±0.04 | 1.13±0.11 | 0.78±0.02 | 1419 | 0 | 0.08 |
|  | Off-phase | 0.49±0.01 | 2.20±0.15 | 1.50±0.18 | 0.52±0.04 | 1.14±0.10 | 0.78±0.02 | 1375 | 0 | 0.04 |
| Probability matching | All | 0.50±0.02 | 1.80±0.13 | 1.55±0.19 | 0.50±0.04 | 0.94±0.08 | 0.84±0.01 | 5792 | 0 | 0.04 |
|  | Control | 0.53±0.01 | 1.42±0.07 | 1.19±0.15 | 0.47±0.04 | 0.81±0.07 | 0.84±0.01 | 3003 | 0 | 0.04 |
|  | On-phase | 0.49±0.02 | 2.11±0.18 | 1.64±0.21 | 0.50±0.04 | 1.05±0.10 | 0.78±0.02 | 1452 | 0 | 0.08 |
|  | Off-phase | 0.49±0.02 | 2.28±0.18 | 1.64±0.21 | 0.50±0.04 | 1.09±0.10 | 0.78±0.02 | 1400 | 0 | 0.15 |
| Stochastic fusion | All | 0.45±0.02 | 2.15±0.19 | 1.92±0.26 | 0.48±0.05 | 0.86±0.05 | 0.76±0.02 | 0 | 0 | 0.08 |
|  | Control | 0.45±0.02 | 1.35±0.09 | 1.48±0.22 | 0.45±0.04 | 0.72±0.05 | 0.74±0.01 | 0 | 0 | 0.08 |
|  | On-phase | 0.43±0.02 | 2.53±0.20 | 1.90±0.25 | 0.49±0.05 | 0.93±0.07 | 0.71±0.02 | 0 | 0 | 0.08 |
|  | Off-phase | 0.44±0.02 | 2.72±0.21 | 1.72±0.19 | 0.48±0.05 | 0.96±0.07 | 0.71±0.02 | 0 | 0 | 0.08 |
| Fixed threshold | All | 1.46±0.09 | 2.21±0.14 | 1.50±0.23 | 0.51±0.04 | 0.97±0.07 | 0.84±0.01 | 5374 | 0 | 0.12 |
|  | Control | 1.45±0.09 | 1.58±0.10 | 1.29±0.22 | 0.49±0.04 | 0.83±0.06 | 0.83±0.01 | 2655 | 0 | 0.04 |
|  | On-phase | 1.42±0.10 | 2.52±0.16 | 1.50±0.21 | 0.51±0.04 | 1.04±0.09 | 0.78±0.02 | 1407 | 0.06 | 0.23 |
|  | Off-phase | 1.45±0.10 | 2.65±0.17 | 1.36±0.15 | 0.51±0.04 | 1.07±0.09 | 0.78±0.02 | 1399 | 0.04 | 0.23 |
| Note: Model parameters (group mean±SEM): pCommon, causal prior; η, probability of stochastic fusion; k, fixed criterion threshold; µ_P_, mean of the numeric prior; σ_P_, standard deviation of the numeric prior; σ_A_, standard deviation of the auditory likelihood; σ_V_, standard deviation of the visual likelihood; Model fit and comparison statistics: R^2^, Nagelkerke’s coefficient of determination^22^ using a null model of random guesses of stimulus number 1-4 with equal probability 0.25; relBIC, Bayesian information criterion at the group level, i.e. subject-specific BICs summed over all subjects (BIC = LL − 0.5 m ln(n), LL = log likelihood, m = number of parameters, n = number of data points) of a model relative to the worst model (n.b. a larger relBIC indicates that a model provides a better explanation of our data); pEP, protected exceedance probability, i.e. the probability that a given model is more likely than any other model, beyond differences due to chance). % win, percentage of participants in which a model won the within-participant model comparison based on BIC. | | | | | | | | | | |

**Table S3**

| **Table S3. Comparison of the BCI model‘s parameters between all pairs of alpha entrainment conditions in the EEG study using paired t-tests.** | | | | | | |
| --- | --- | --- | --- | --- | --- | --- |
| **BCI parameter** | **Alpha entrainment** | **t** | **p** | **p_corr_** | **Cohen‘s d** | **BF_10_** |
|  | Con vs. On | 2.883 | 0.008 | 0.074 | 0.565 | 5.705 |
| pCommon | Con vs. Off | 2.898 | 0.006 | 0.064 | 0.568 | 5.889 |
|  | On vs. Off | 0.140 | 0.889 | 1 | 0.027 | 0.209 |
|  | Con vs. On | -4.454 | <0.001 | <0.001 | -0.874 | 181.783 |
| µ_P_ | Con vs. Off | -4.884 | <0.001 | <0.001 | -0.958 | 495.462 |
|  | On vs. Off | -1.744 | 0.068 | 0.271 | -0.342 | 0.778 |
|  | Con vs. On | -2.628 | 0.004 | 0.048 | -0.515 | 3.461 |
| σ_P_ | Con vs. Off | -2.556 | 0.009 | 0.074 | -0.501 | 3.021 |
|  | On vs. Off | -0.100 | 0.929 | 1 | -0.020 | 0.208 |
|  | Con vs. On | -2.282 | 0.028 | 0.196 | -0.448 | 1.839 |
| σ_A_ | Con vs. Off | -1.981 | 0.061 | 0.293 | -0.389 | 1.114 |
|  | On vs. Off | 0.659 | 0.555 | 1 | 0.129 | 0.253 |
|  | Con vs. On | -4.130 | <0.001 | <0.001 | -0.810 | 86.004 |
| σ_V_ | Con vs. Off | -4.558 | <0.001 | <0.001 | -0.894 | 231.399 |
|  | On vs. Off | -2.036 | 0.052 | 0.293 | -0.399 | 1.215 |
| Note: The BCI model’s decision strategy applies model averaging. BCI Model parameters: pCommon, causal prior; η, probability of stochastic fusion; k, fixed criterion threshold; µ_P_, mean of the numeric prior; σ_P_, standard deviation of the numeric prior; σ_A_, standard deviation of the auditory likelihood; σ_V_, standard deviation of the visual likelihood; Alpha entrainment: control, Con; on-phase, On; off-phase, Off. p values were derived from a paired two-sided randomisation test based on a t statistic (n = 5000 randomisations); p_corr_ is multiple-comparison corrected across the 15 comparisons using the Bonferroni-Holm correction. Bayes factors BF_10_ were computed in Bayesian t-tests assuming a Jeffrey-Zellner-Siow prior and a scaling factor of 0.707 for the Cauchy prior. | | | | | | |

**Table S4**

| **Table S4. Results of the Bayesian model comparison in the psychophysical study. The BCI model with three decision strategies (model averaging MA, model selection MS and probability matching PM) and heuristic models (stochastic fusion SF and fixed threshold FT) were compared, separately for the three different entrainment frequencies (EF).** | | | | | | | | | | | | | | | | | | | | | | | | | |
| --- | --- | --- | --- | --- | --- | --- | --- | --- | --- | --- | --- | --- | --- | --- | --- | --- | --- | --- | --- | --- | --- | --- | --- | --- | --- |
| **EF** | **Mo-del** | **pCommon**  **η**  **k** | | | | | | | | | **µ_P_** | | **σ_P_** | | **σ_A_** | | **σ_V_** | | ***R*^2^** | | **rel**  **BIC** | | **pEP** | | **% win** |
|  |  | **0π/4** | **1π/4** | **2π/4** | **3π/4** | **4π/4** | **5π/4** | **6π/4** | **7π/4** |  | |  | |  | |  | |  | |  | |  | |  | |
| 5 Hz | MA | 0.55 | 0.55 | 0.55 | 0.53 | 0.53 | 0.54 | 0.55 | 0.54 | 1.63 | | 1.00 | | 0.35 | | 0.89 | | 0.80 | | 2483 | | | 1.00 | 0.79 | |
|  | MS | 0.55 | 0.56 | 0.56 | 0.54 | 0.54 | 0.55 | 0.55 | 0.55 | 1.79 | | 0.83 | | 0.36 | | 0.88 | | 0.79 | | 2101 | | | 0.00 | 0.07 | |
|  | PM | 0.57 | 0.57 | 0.57 | 0.55 | 0.55 | 0.56 | 0.57 | 0.56 | 1.76 | | 0.84 | | 0.34 | | 0.82 | | 0.79 | | 2116 | | | 0.00 | 0.07 | |
|  | SF | 0.42 | 0.42 | 0.42 | 0.42 | 0.41 | 0.43 | 0.43 | 0.43 | 1.69 | | 0.98 | | 0.32 | | 0.75 | | 0.72 | | 0 | | | 0.00 | 0.00 | |
|  | FT | 1.32 | 1.31 | 1.28 | 1.26 | 1.26 | 1.32 | 1.31 | 1.34 | 1.80 | | 0.82 | | 0.36 | | 0.87 | | 0.78 | | 1847 | | | 0.00 | 0.07 | |
| 10 Hz | MA | 0.55 | 0.54 | 0.54 | 0.53 | 0.54 | 0.54 | 0.54 | 0.55 | 1.57 | | 1.05 | | 0.35 | | 0.90 | | 0.80 | | 2687 | | | 1.00 | 0.76 | |
|  | MS | 0.56 | 0.55 | 0.55 | 0.54 | 0.55 | 0.56 | 0.55 | 0.56 | 1.77 | | 0.84 | | 0.36 | | 0.89 | | 0.79 | | 2329 | | | 0.00 | 0.03 | |
|  | PM | 0.57 | 0.56 | 0.56 | 0.55 | 0.56 | 0.57 | 0.56 | 0.57 | 1.73 | | 0.86 | | 0.34 | | 0.81 | | 0.79 | | 2350 | | | 0.00 | 0.07 | |
|  | SF | 0.44 | 0.40 | 0.43 | 0.40 | 0.44 | 0.43 | 0.41 | 0.44 | 1.65 | | 0.99 | | 0.38 | | 0.86 | | 0.70 | | 0 | | | 0.00 | 0.03 | |
|  | FT | 1.34 | 1.30 | 1.33 | 1.26 | 1.33 | 1.34 | 1.31 | 1.35 | 1.74 | | 0.83 | | 0.36 | | 0.85 | | 0.78 | | 2145 | | | 0.00 | 0.10 | |
| 15 Hz | MA | 0.53 | 0.53 | 0.53 | 0.54 | 0.54 | 0.53 | 0.54 | 0.55 | 1.53 | | 1.03 | | 0.34 | | 0.91 | | 0.80 | | 2431 | | | 1.00 | 0.83 | |
|  | MS | 0.54 | 0.54 | 0.54 | 0.54 | 0.56 | 0.54 | 0.54 | 0.56 | 1.73 | | 0.83 | | 0.36 | | 0.89 | | 0.79 | | 2009 | | | 0.00 | 0.07 | |
|  | PM | 0.55 | 0.55 | 0.55 | 0.55 | 0.56 | 0.55 | 0.55 | 0.57 | 1.69 | | 0.86 | | 0.34 | | 0.82 | | 0.79 | | 2040 | | | 0.00 | 0.03 | |
|  | SF | 0.41 | 0.41 | 0.42 | 0.42 | 0.44 | 0.41 | 0.42 | 0.43 | 1.54 | | 0.95 | | 0.31 | | 0.71 | | 0.72 | | 0 | | | 0.00 | 0.00 | |
|  | FT | 1.27 | 1.29 | 1.30 | 1.27 | 1.34 | 1.26 | 1.28 | 1.32 | 1.63 | | 0.81 | | 0.35 | | 0.84 | | 0.79 | | 1856 | | | 0.00 | 0.07 | |
| Note: Model parameters (group mean): pCommon, causal prior; η, probability of stochastic fusion; k, fixed criterion threshold; µ_P_, mean of the numeric prior; σ_P_, standard deviation of the numeric prior; σ_A_, standard deviation of the auditory likelihood; σ_V_, standard deviation of the visual likelihood; Model fit and comparison statistics: R^2^, Nagelkerke’s coefficient of determination^22^ using a null model of random guesses of stimulus number 1-4 with equal probability 0.25; relBIC, Bayesian information criterion at the group level, i.e. subject-specific BICs summed over all subjects (BIC = LL − 0.5 m ln(n), LL = log likelihood, m = number of parameters, n = number of data points) of a model relative to the worst model (n.b. a larger relBIC indicates that a model provides a better explanation of our data); pEP, protected exceedance probability, i.e. the probability that a given model is more likely than any other model, beyond differences due to chance). % win, percentage of participants in which a model won the within-participant model comparison based on BIC. | | | | | | | | | | | | | | | | | | | | | | | | | |
